# How to start a LINE: 5’ switching rejuvenates LINE retrotransposons in tobacco and related *Nicotiana* species

**DOI:** 10.1101/2022.11.04.512097

**Authors:** Nora Hartig, Kathrin M. Seibt, Tony Heitkam

## Abstract

In contrast to their conserved mammalian counterparts, plant long interspersed nuclear elements (LINEs) are highly variable, splitting into many low-copy families. Curiously, LINE families from the RTE clade retain a stronger sequence conservation and hence reach higher copy numbers. The cause of this RTE-typical property is not yet understood, but would help clarifying why some transposable elements are removed quickly whereas others persist in plant genomes. Here, we bring forward the first detailed study of RTE LINE structure, diversity and evolution in plants. For this, we argue that the Nightshade family is the ideal taxon to follow the evolutionary trajectories of RTE LINEs, given their high abundance, recent activity and partnership to non-autonomous elements.

Using bioinformatic, cytogenetic and molecular approaches, we detect 4029 full-length RTE LINEs across the *Solanaceae*. We finely characterize and manually curate a core group of 458 full-length LINEs in allotetraploid tobacco, show amplification after polyploidization, and trace hybridization events by RTE LINE composition of parental genomes. Finally, we reveal the role of the untranslated regions (UTRs) as causes for the unique RTE LINE amplification and evolution pattern in plants: On one hand, we detect a highly conserved motif at the 3’ UTR, suggesting strong selective constraints acting on the RTE terminus. On the other hand, we observed successive rounds of 5’ UTR cycling, constantly rejuvenating the promoter sequences. This interplay between exchangeable promoters and conserved LINE bodies and 3’ UTR likely allows RTE LINEs to persist and thrive in plant genomes.

## Introduction

Although strikingly abundant in higher mammals, long interspersed nuclear elements (LINEs) are among the least proliferating retrotransposons in flowering plants (Kumar et al. 1999; Renny-Byfield et al. 2011; Bombarely et al. 2016; Ivancevic et al. 2016; Sierro et al. 2018). Nevertheless, plant LINEs carry the important key enzymes – a reverse transcriptase (RT) and an endonuclease (EN) – to produce numerous copies in a plant host. For this, the LINE-encoded machinery nicks the genomic host DNA and integrates a retro-copy using the target-primed reverse transcription mechanism (Luan et al. 1993; Ichiyanagi and Okada 2008; Kapitonov et al. 2009). In consequence, LINE amplification may increase genome sizes (Lander et al. 2001; Gentles et al. 2007), influence genome structure and dynamics (Belancio et al. 2008), and may lead to species diversification (Guichard et al. 2018).

For their proliferation, LINE expression and subsequent reverse transcription are mandatory. Expression is controlled by an internal RNA polymerase II promoter located at the LINE head, the 5’ untranslated region (UTR). On the opposing LINE end, the 3’ UTR harbours the recognition sequence for the LINE-encoded RT, thus priming the reverse transcription into cDNA (Swergold 1990; Luan et al. 1993; Smit et al. 1995; Okada et al. 1997; Kapitonov et al. 2009; Hayashi et al. 2014). Hence, differences within the UTRs are crucial to selectively control LINE populations within a genome. This UTR-driven control of LINE amplification has been suggested as a mammalian-specific phenomenon (Adey et al. 1994; Khan et al. 2006; Sookdeo et al. 2018).

In plants, the “LINE-1” (L1) and “retrotransposable element” (RTE) LINEs constitute the most prominent LINE clades (Heitkam et al. 2014), and belong to the most ancient retrotransposon lineages (Župunski et al. 2001). Plant L1 LINEs are generally much more variable in sequence and structure than their mammalian counterparts. Hence, they split into many L1 families with only very few members (Heitkam et al. 2014). In contrast, RTE LINEs are generally much more conserved, hence forming only few families with many members (Heitkam et al. 2014). It is not yet understood, why these two LINE types evolve in fundamentally different manners, and main cause of this is the lack of knowledge on plant RTE LINEs.

Structurally, members of the RTE-clade LINEs are distinguished from other LINE types by the presence of a single, continuous ORF and the absence of a dedicated *gag*-like region. They are often terminated by a microsatellite tail, typically a [TTG]_n_ motif, instead of the canonical poly-(A)-tail described for L1 LINEs (Malik and Eickbush 1998; Wicker et al. 2007). Recent reports show that RTE LINEs are widespread in the angiosperms (Ivancevic et al. 2016, 2018; Gao et al. 2018), with a considerable sequence conservation in their coding regions (Heitkam et al. 2014).

We argue that the *Solanaceae* (Nightshades) may represent an ideal framework for the investigation of RTE LINE biology in plant genomes: First, these genomes harbor the SolRTEs, an RTE LINE group marked by stucturally similar open reading frames and low genetic divergence (Wenke et al. 2011; Heitkam et al. 2014), potentially indicating recent mobilization. Second, RTE LINEs are widely distributed among this plant family (Renny-Byfield et al. 2011; Sierro et al. 2013, 2018; Bombarely et al. 2016; Ivancevic et al. 2018), allowing insights into RTE LINE evolution. Third, some of these SolRTEs are transcribed, associated with genic regions, and may impact metabolism regulation (Wenke et al. 2011; Mehra et al. 2015; Jung et al. 2019). Finally, although often hypothesized (Okada et al. 1997; Kajikawa and Okada 2002; Gogolevsky et al. 2008), sequence-dependent partnerships between long and short interspersed nuclear elements (LINEs and SINEs) have very rarely been observed in plants. In such a partnership, SINEs may utilize the enzymatic machinery of their autonomous partner LINE, based on a shared and highly similar region of the 3’ UTR (Ohshima et al. 1996; Okada et al. 1997; Piskurek et al. 2006). Especially in tobacco (*Nicotiana tabacum*), the SolRTEs are widely-accepted as autonomous and sequence-dependent partner LINEs to a non-autonomous SINE family, the TS-SINEs (Yoshioka et al. 1993; Ohshima et al. 1996; Wenke et al. 2011). As recent TS-SINE amplification was reported for *N. tabacum* (Mhiri et al. 2019), this also implies the recent activity SolRTE LINEs in this genome. Thus, taking all points together, SolRTEs are optimally suited to investigate the impact of RTE LINE amplification on plant genomes. To provide a baseline for the understanding of RTE biology, we aimed to trace their evolution across the Nightshades, focussing especially on their contribution to the allotetraploid *N. tabacum* genome.

*N. tabacum* is one of 76 naturally occuring species in the genus *Nicotiana* within the *Solanaceae* plant family. As nearly half of these species are allopolyploid (Knapp et al. 2004), this genus is marked by the ability to combine several parental genome sets within a single nucleus. These allopolyploidization events may induce genomic shocks (McClintock 1984), likely the result of an imbalance of parental transposable elements (Lim et al. 2007; Michalak 2009; Freeling et al. 2012; Parisod et al. 2012). Tobacco belongs to the most frequently studied species of this genus (e.g. Clarkson et al. 2005, 2017; Lim et al. 2007). It represents one of the youngest allotetraploids within the genus (Clarkson et al. 2005), harboring the highly diverged subgenomes of *N. tomentosiformis* and *N. sylvestris*. In synthetic *N. tabacum* allopolyploids, transposable element activity was already observed (Mhiri et al. 2019), supporting the genomic shock hypothesis.

Benefitting from the SolRTE’s widespread abundance and their presumed roles in transposable element control (e.g. control of SINEs) within the *Nicotiana* genus, here, we focus on this plant group to shed light into the evolution and biological roles of the SolRTEs. Thus, we follow their evolution on a stuctural and sequence level across the speciation events within the *Nicotiana* genus, especially focusing on the allopolyploidization of *N. tabacum*. To provide a robust basis for all downstream analyses, we identify and finely classify the *N. tabacum* SolRTE population and follow their distribution across the genus *Nicotiana* and further *Solanaceae* species. Using a combination of bioinformatic approaches and molecular cytogenetics, we investigate the role of the UTRs for RTE LINE amplification and evolution. On one hand, we identify a highly conserved structure within their 3’ UTRs, suggesting strong selective constraints and potentially mechanistic impacts. On the other hand, we observe several rounds of UTR cycling, accompanied by the formation of new SolRTE lineages. To integrate our observations, we discuss a possible scenario for the formation of parent-specific SolRTE lineages in the subgenomes of *N. tabacum*. Due to their typical lineage-specific UTRs, these newly arisen SolRTE lineages can be traced in a parent-depending way, allowing to follow hybridization and chromosomal restructuring events in the young polyploid tobacco genome.

## Results

### Tobacco harbours 458 full-lengths SolRTE LINEs that form four families, eleven subfamilies and nine variants

To capture the full structural diversity of RTE LINEs in *N. tabacum*, we aimed to identify all full-length members. For this, we created nHMMs for the specific detection of RTs and ENs of the RTE type. Multiple filtering steps were applied to exclude highly diverged and truncated RTE candidates as well as putative assembly errors (Notes S1). Although a particularly high portion of candidates (n = 3247) was 5‘ truncated we finally extracted 458 full-length SolRTE members from the genome of *N. tabacum*. This robust set of elements builds the foundation for the structural characterization and downstream analyses.

The ORFs of these 458 full-length elements are very conserved (Figure 1A), sharing a nucleotide identity of 73.2 %. In contrast, the UTRs are more variable. While the 5’ UTR differs in length and sequence, the 3’ UTR exhibited a bipartite nature. It has a variable (v3’ UTR) and conserved part (c3’ UTR), arranged sequentially in upstream direction. All c3’ UTRs share an identity of 68.6 % over all analyzed copies. RTE elements were 3’ terminated by the characteristic [TTG]_n_ tail.

**Figure 1:**
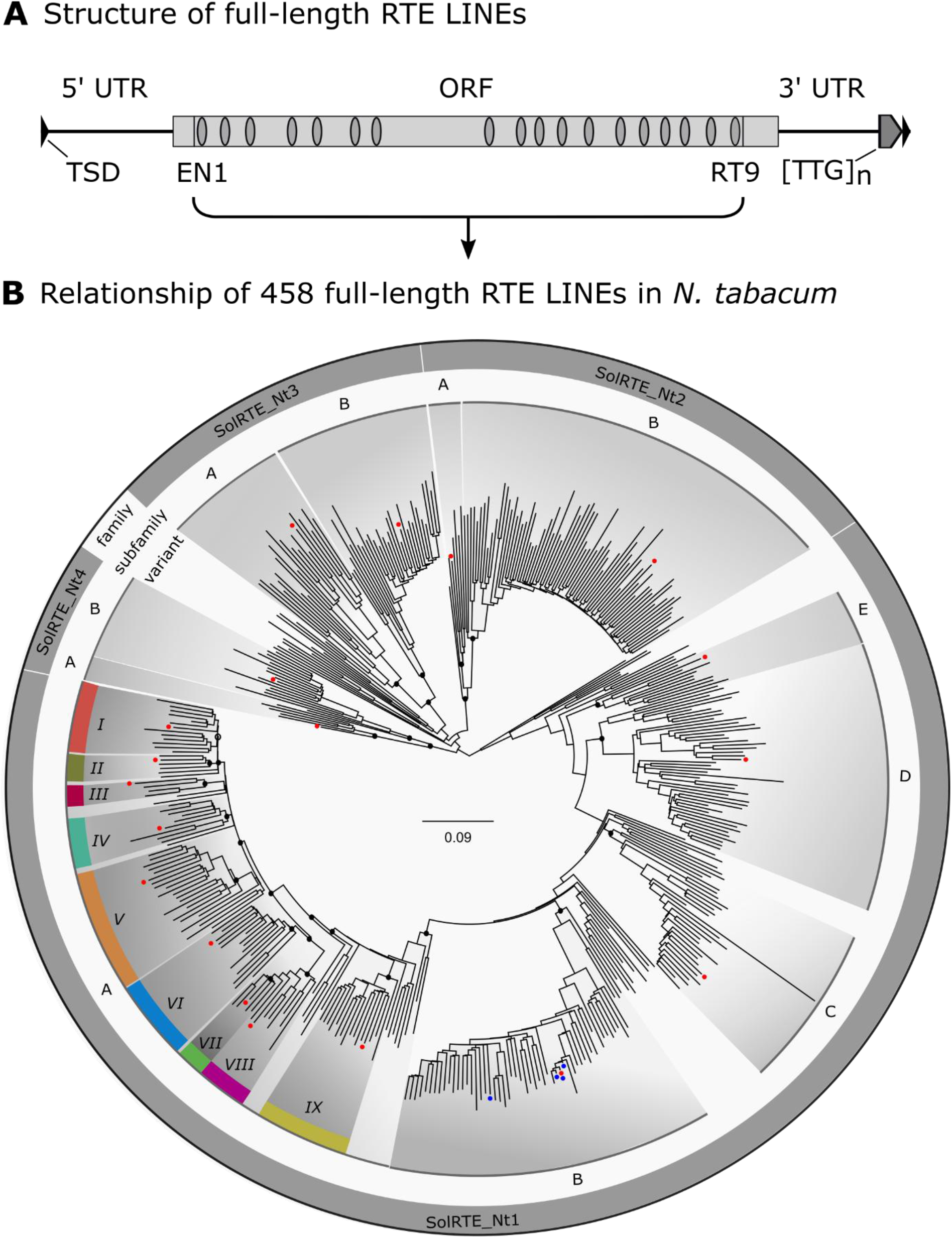
**(A) Schematic structure of a typical RTE LINE in *Nicotiana tabacum*.** Rectangles depict open reading frames (ORFs), whereas triangles show target site duplications (TSDs). Conserved apurinic endonuclease (EN) and reverse transcriptase (RT) protein domains according to Wenke *et al*. (2011) are indicated by ovals. The repetitive [TTG]_n_ motif terminating the RTE LINE is marked. **(B) Relationships between the 458 RTEs that were identified from the *N. tabacum* ‘TN90’ assembly**. The phylogram was calculated using the Maximum Likelihood method and was based on the conserved nucleotide sequences (4447 bp) that span the ORF from *EN1* up to *RT9*. Bootstrap node support values were obtained from 1000 resamples. Nodes of interest were marked with black dots, if support values were >95, or black circles with node support >60. Representative copies of each family, subfamily and variant were marked with red dots within the phylogram. RTE copies with complete and continuous ORFs were marked with blue dots.

As the variability of 5’ UTR and v3’ UTR suggested a diversity among the 458 tobacco SolRTEs, we used cluster and maximum likelihood (ML) analyses to classify the RTE LINE population of *N. tabacum*. Based on the multiple sequence alignment of the ORF (458 sequences, 4447 bp), our k-means cluster analyses suggested a separation into five main clusters (Figure S1), named SolRTE_Nt1 to SolRTE_Nt5, with a mean genetic distance ranging from 0.18 to 0.31 within one cluster (Figure S1C). To test their monophyletic origin, we compared the identified clusters to the ML tree. The RaxML analysis supported the monophyly of only four of the five clusters (SolRTE_Nt1 to SolRTE_Nt4), whereas the remaining SolRTE_Nt5 sequence group was deemed paraphyletic (Figure 1B). As this cluster also showed a high intra-genetic distance of 31 % (Figure S1C), with unclear phylogenetic relationship and classification, SolRTE_Nt5 was excluded from further analyses. Based on the monophyletic character of the clusters SolRTE_Nt1 to SolRTE_Nt4, we classified these as distinct SolRTE families.

Comparison of inter- and intra-family distances of the four SolRTE families in *N. tabacum* revealed further diversification within each sequence group (Figure S1 & S2). Thus, we subdivided the four SolRTE_Nt families into eleven subfamilies, named by appending a capital Latin letter to the group name (e.g. subfamily SolRTE_Nt1A-E).

All SolRTEs within a subfamily are marked by a shared and subfamily-specific v3’ UTR. The subfamily members are highly similar to each other starting from the ORF up to the tail. However, following up the SolRTE_Nt LINE in the upstream direction, we observed a decrease in similarity within the 5’ UTR, and therefore, a tremendous increase in the number of variants, with some copies being unique regarding their 5’ UTR. As SolRTE_Nt1A represents the biggest SolRTE subfamily within the tobacco genome, it was exemplarily chosen for manual subdivision into variants based on the 5’ UTRs. Nine SolRTE_Nt1A variants were identified and named by appending a roman number for their specific, non-homologous 5’ UTR variant. In addition to k-means and hierarchical clustering, SiLiX cluster analysis (Figure S3) further supports the subfamily structure established here.

To identify structural and sequence characteristics among tobacco RTE LINEs, reference elements for each subfamily and variant were chosen and deeply characterized (Figure 2, Table S1). All members of the SolRTE-Nt1A subfamily showed a highly similar v3’ UTR, except for the variants *VII* and *IIX*. Both variants show the same deletion of 73 bp and 76 bp compared to variant *IX*, respectively. Within the otherwise non-homologous and variant-specific 5’ UTR sequences, the variants *VI* and *VII* share two highly identical regions, with 73.8 % identity over 145 bp and 81 % identity over 105 bp.

**Figure 2:**
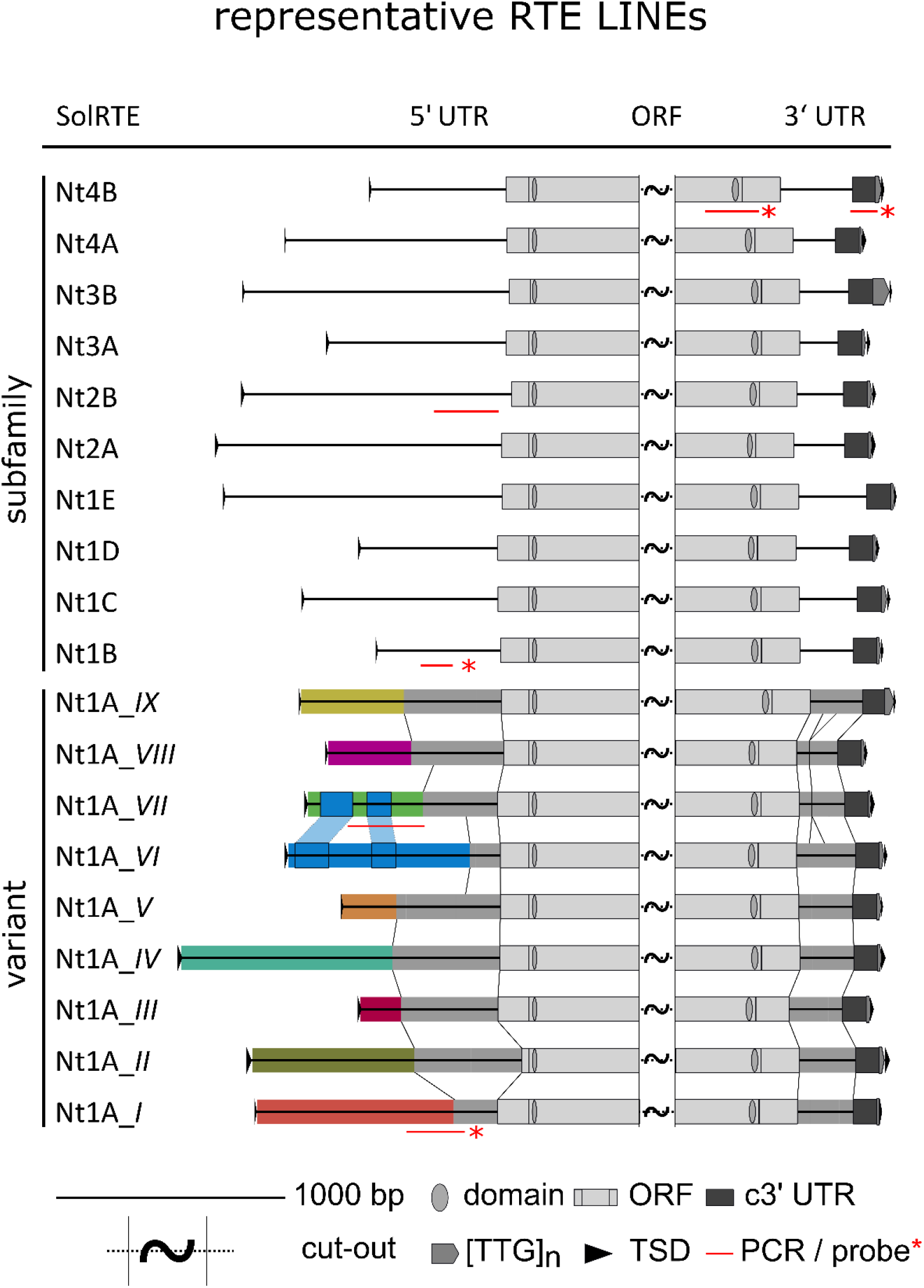
Structure of 19 representative RTE LINEs, illustrating their structural diversity in *N. tabacum*. The depicted retroelements correspond to each SolRTE family (SolRTE_Nt1 to SolRTE_Nt4) and subfamily (marked by the appendices A-E). The largest subfamily, SolRTE_Nt1A, was further divided into nine variants (SolRTE_Nt1A_*I* to SolRTE_ Nt1A_*IX*). Grey rectangles in the 5’ and 3’ UTRs show shared sequences among different variants, whereas differently colored regions show non-homologous parts of the 5’ UTR. For simplicity middle part of the open reading frame (ORF) was cropped (wave). The narrow red lines depict the regions used for the experiments in Figure 3, 5 and 6. Abbreviations are as follows: Open reading frame (ORF), untranslated region (UTR), conserved 3’ UTR (c3’ UTR), repetitive motif at the 3’ terminus ([TTG]_n_), target site duplication (TSD).

We identified putative promoter motifs (e.g. numerous TATA boxes), common *cis*-acting elements in promoter and enhancer regions (CAAT – box) and putative DNA motifs related to light, abiotic stress, and plant hormones. Less frequent motifs were associated with defense and stress, protein binding sites, endosperm and meristem expression, cell cycle regulation and circadian control with presence in only some regions.

### RTE LINEs are widespread and structurally conserved in the genus *Nicotiana* and other Nightshades

To explore the abundance of SolRTEs within the genus *Nicotiana* and other members of the *Solanaceae* plant family, we retrieved all available *Nicotiana* and five additional Nightshade genome assemblies (Table S2). Using the nHMM strategy as outlined above, these genomes were analysed bioinformatically under the same conditions as *N. tabacum*. SolRTEs were found to occur in all analysed Nightshade species, with the SolRTE copy number mainly depending on the genome size (Figure S4 & Table S2). Using hierarchical clustering, SolRTEs of each genus were separated according to the genetic distance of their ORFs (Figure 3A), with *Nicotiana, Solanum, Capsicum*, and *Petunia* SolRTEs forming distinct clusters, each. SolRTEs extracted for *N. tabacum* were not exclusively assigned into one separated cluster, i.e. they were distributed over subclusters containing SolRTEs of different *Nicotiana* species, indicating the identified subfamilies and variants also being present in other *Nicotiana* species.

**Figure 3:**
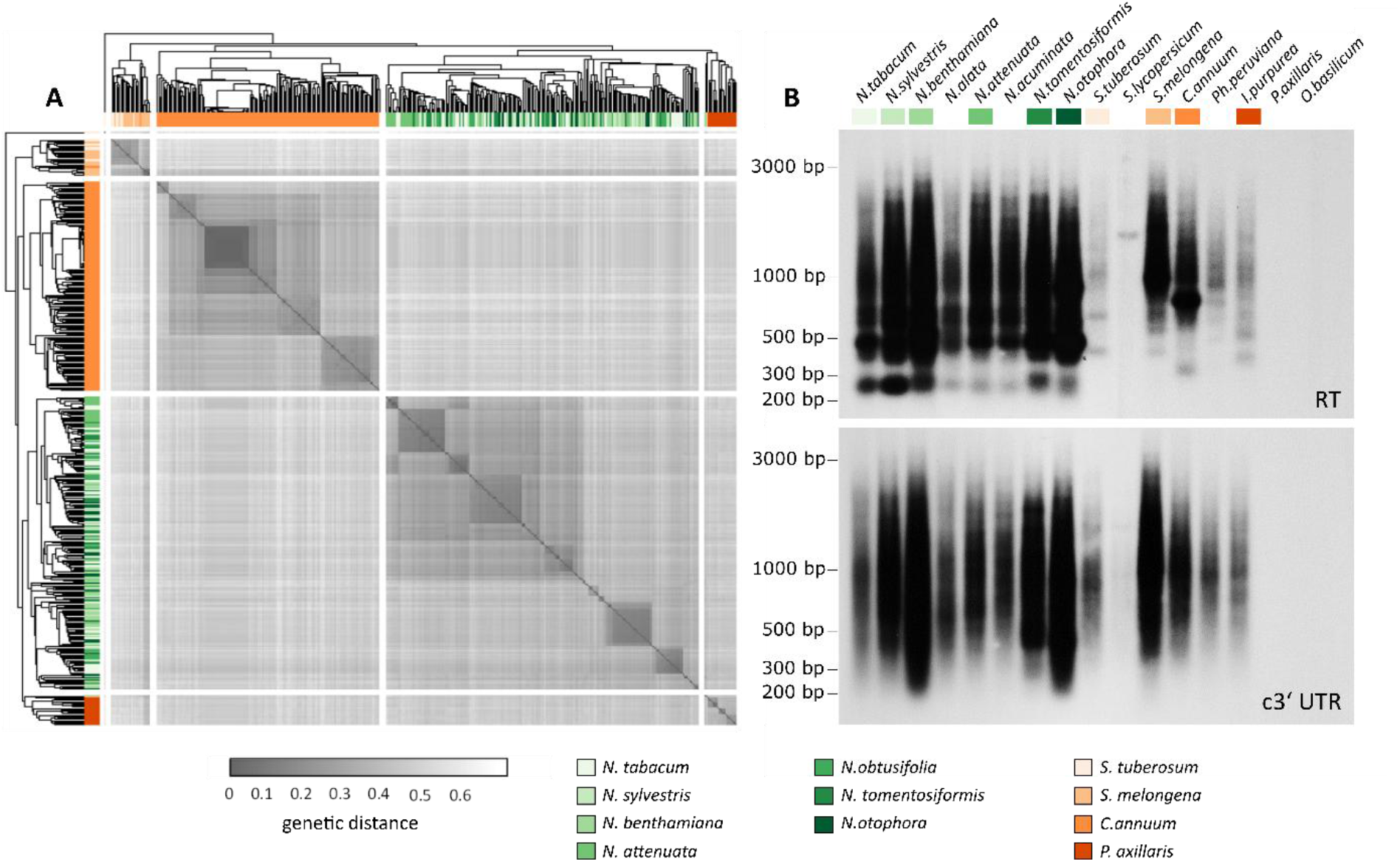
RTE LINEs are abundant and structurally conserved across the plant family *Solanaceae*. (**A)** Heatmap of genetic distances between 420 RTE LINEs from eleven species representative of SolRTE diversity within *Solanaceae*. Colors indicate the affiliation of each randomly selected SolRTE copy to the respective species analyzed (green shades for *Nicotiana* species, orange shades for other nightshade species). Using the TN93 substitution model (Tamura and Nei, 1993), genetic distances were calculated for the conserved nucleotide sequences spanning the open reading frame from the first apurinic endonuclease to the last reverse transcriptase protein domains.The intensity of the grey shading corresponds to the genetic distance. The dendrogram and clustering are based on UPGMA distance method (Sokal and Michener, 1958). **(B)** Comparative autoradiograms of Southern hybridizations with RTE-derived probes onto *Hinf*I-restricted genomic DNA. Species from the nightshades (including the genera *Nicotiana, Solanum, Capsicum*, and *Petunia*) and three outgroups (*Ipomoeae purpurea, Ocimum basilicum, Beta vulgaris*) were included. Conserved regions of the reverse transcriptase (RT) and the conserved 3’ UTR (c3’ UTR) were amplified from *N. tabacum* and used as probes. Autoradiograms were exposed for 18 h (RT) and for 72 h (c3’ UTR).

To verify the SolRTE distribution detected bioinformatically in the *Solanaceae* reference genomes, we used Southern hybridisation to explore the presence of SolRTEs in 14 Nightshade species and three species outside the plant family (*I. purpurea, O. basilicum, B. vulgaris*) using probes of RT domain 9 and c3’ UTR (Figure 3B). Both structural components are highly conserved within the Nightshade RTEs and with only few exemptions they show strong hybridisation signals on the autoradiograms (Figure 3B, lanes 1-14). In contrast, no signals were detected in the three outgroup species (lanes 15-17). The nearly identical signal patterns for RTs suggest a sequence conservation within the genus *Nicotiana* (lanes 1-8) and differentiation to other analyzed species (9-14). The length of the observed RT restriction fragments correspond to the bioinformatically predicted sizes for *N. tabacum* (e.g. approximately 250 bp, 470 bp).

To summarize, RTE-LINEs of the SolRTE-type are distributed over the whole plant family *Solanaceae*, but not in any of the analysed outgroup species, even not in the next closely related family *Convolvulaceae* (*I. purpurea*). Although RTE LINEs of each genus were diverged from each other, all RTE LINEs within the plant family *Solanaceae* have a conserved sequence in their c3’ UTR. The abundance of RTEs among Nightshades varies, with a comparatively high number of copies within the members of the *Nicotiana* genus. The identified families for *N. tabacum* (SolRTE_Nt1-4) are also present in other *Nicotiana* species.

### The SolRTE body is conserved across the *Nicotiana*, with species- and subgenome-specific variants

As clustering of the SolRTE sequences based on their genetic distances revealed a wide distribution of different variants over the genus *Nicotiana*, we analyzed the species-specific genomic RTE landscapes in detail (Figure 4 & 5).

**Figure 4:**
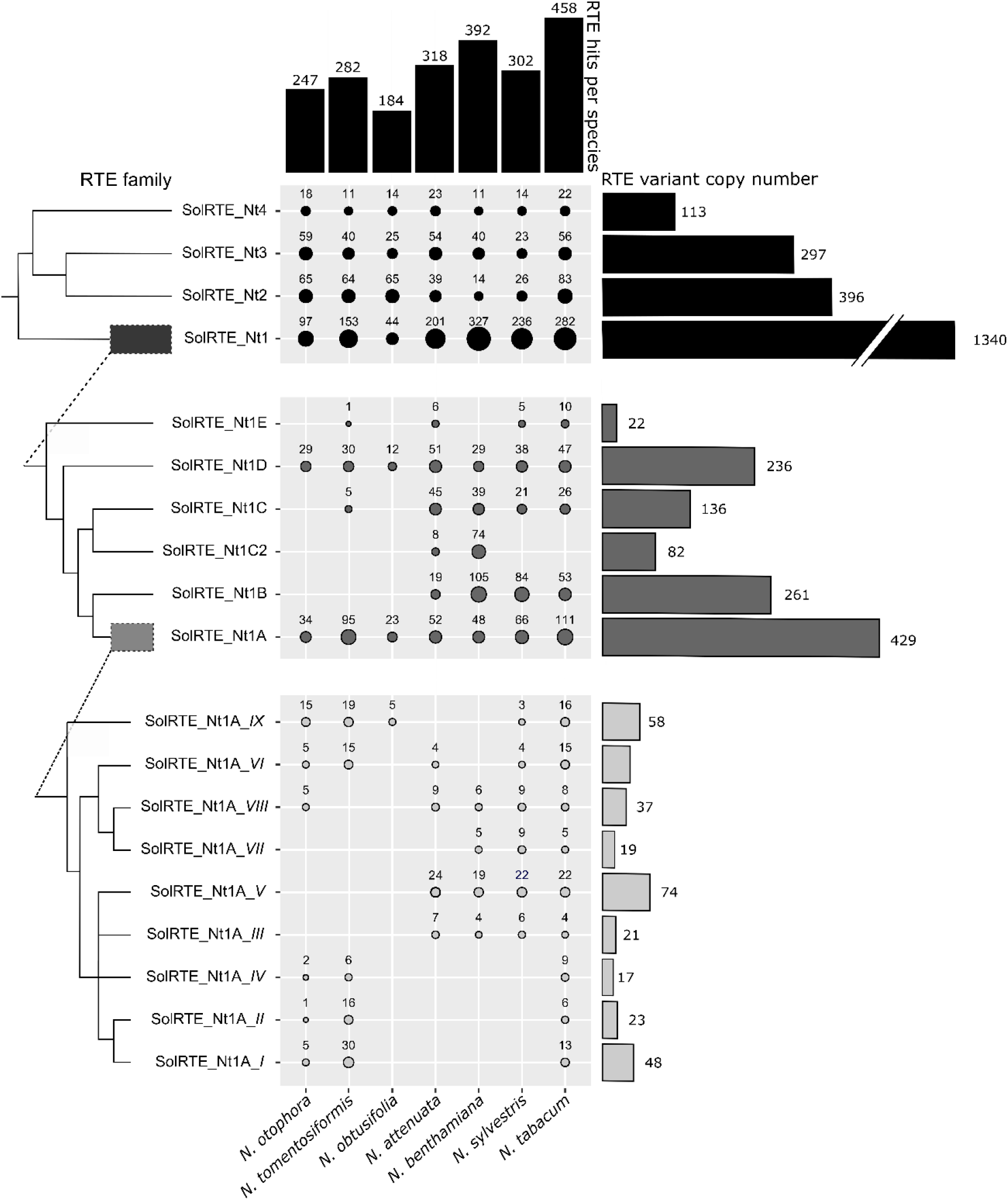
Distribution and abundance of RTE LINEs within the genus *Nicotiana*. The bubble chart shows the contribution of each identified SolRTE-family (Nt1-4), subfamily (marked by the appendices A-E) and the nine variants of the subfamily SolRTE_Nt1A (SolRTE_Nt1A_*I* – *IX*) to RTE LINE diversity in the genomes of seven *Nicotiana* species. Circle areas correlate with the number of identified full length elements. For comparison, the bars at the top summarize the number of full length elements per genome, while the bars on the right summarize the number of full length elements of each SolRTE family, subfamily and variant across all the species analyzed. The dendrogram on the left represents the relationship among the classified SolRTE family, subfamily and variants, based on the phylogram from Figure 1B, calculated using the Maximum Likelihood method.

**Figure 5:**
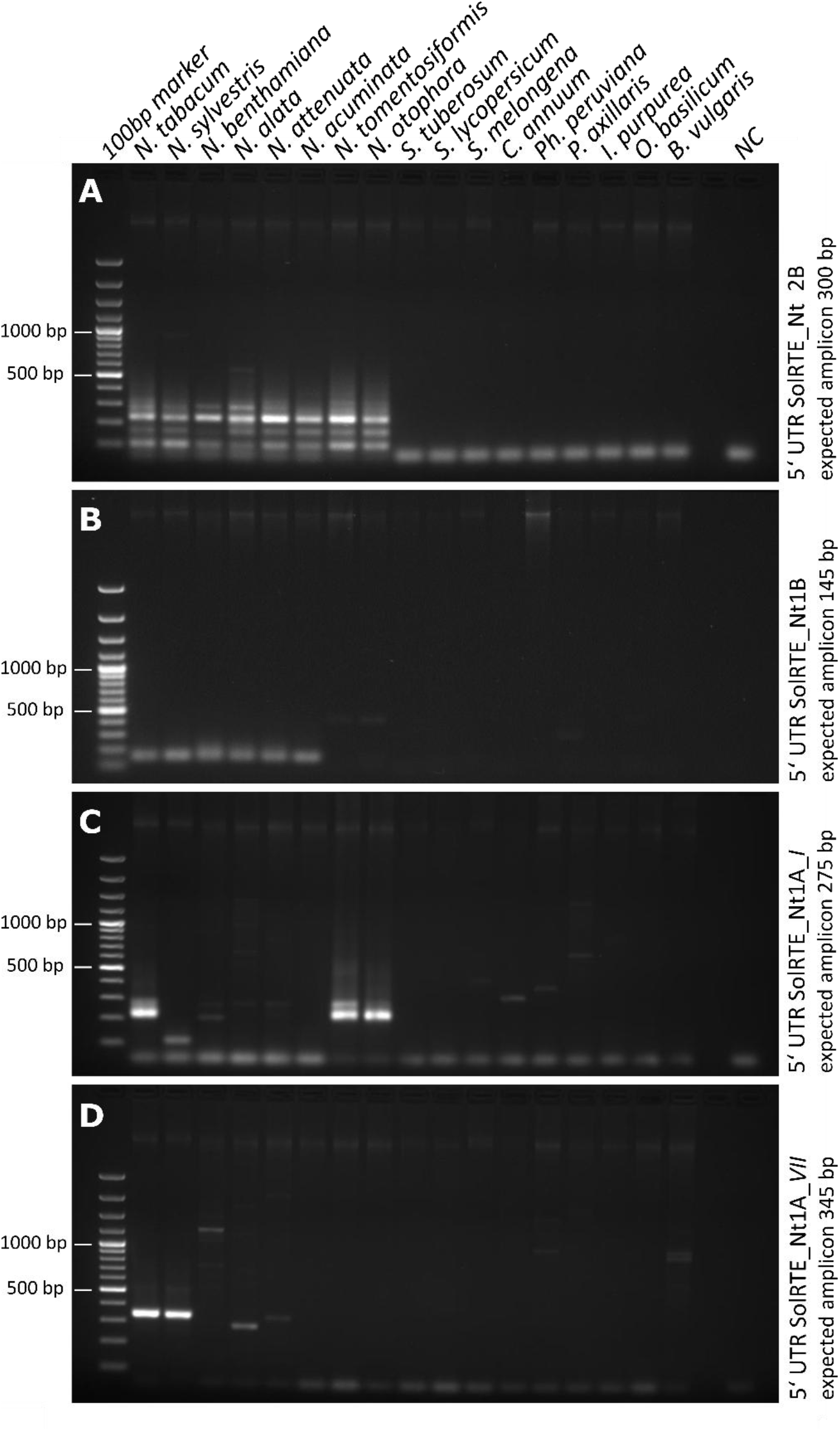
Comparative PCRs distinguishing species-specific SolRTE variants. All primer pairs amplify LINE 5’ UTRs (as indicated in Figure 2) of the different subfamilies SolRTE_Nt2B **(A)** and SolRTE_Nt1B **(B)**, as well as the variants SolRTE_Nt1A_*I* **(C)** and SolRTE_Nt1A_*VII* **(D)** within the plant family *Solanaceae* (*Nicotiana, Solanum, Capsicum, Petunia*) and three outgroup species (*Ipomoeae purpurea, Ocimum basilicum, Beta vulgaris*). Expected amplicon sizes are stated on the right.

The families SolRTE_Nt2 to 4 and subfamilies SolRTE_Nt1C to E are distributed over all seven analyzed *Nicotiana* species (*N. attenuata*, *N. benthamiana*, *N. otophora*, *N. sylvestris*, *N. tabacum*, *N. tomentosiformis*). In contrast other subfamilies or variants are only detected within certain species, namely SolRTE_Nt1C2 for *N. benthamiana* and *N. attenuata;* SolRTE_Nt1B for *N. benthamiana N. attenuata, N. sylvestris*, and *N. tabacum;* SolRTE_Nt1A*_ VII* for *N. benthamiana, N. sylvestris*, and *N. tabacum;* as well as SolRTE_Nt1A_*I* for *N. tomentosiformis, N. otophora*, and *N. tabacum* (Figure 4).

To validate the presence of different subfamilies and variants in the *Nicotiana* genomes, comparative PCR analyses with 14 Nightshade species and three species outside the plant family (*I. purpurea, O. basilicum, B. vulgaris*) were conducted. The specific regions of the 5’ UTRs were amplified for four differently distributed SolRTE subfamilies and variants (Figure 5): (1) the subfamily SolRTE_Nt2B is distributed throughout the genus *Nicotiana* (Figure 5A, lanes 2 - 9), (2) the subfamily SolRTE_Nt1B is *Nicotiana*-specific (Figure 5B, lanes 2 - 7), excluding the diploid species *N. tomentosiformis and N. obtusifolia* (Figure 5B, lanes 8 - 9), (3) the variant SolRTE_Nt1A_*I* is specific for the diploids *N. tomentosiformis* and *N. otophora*, and in tetraploid *N. tabacum* genomes (Figure 5C, lanes 9, 8 and 2, respectively), and (4) the variant SolRTE_Nt1A_*VII* occurs only in diploid *N. sylvestris* and in the tetraploids *N. tabacum* (Figure 5D, lanes 3 and 2, respectively).

Taking into account the hybrid origin of allopolyploid *N. tabacum*, the subfamily SolRTE_Nt1B and the variants SolRTE_Nt1A_*III*, SolRTE_Nt1A_*V* and SolRTE_Nt1A_*IIV* were detected only within the genome of the maternal species *N. sylvestris*, whereas the variants SolRTE_Nt1A_*I*, SolRTE_Nt1A_*II*, and SolRTE_Nt1A_*IV* were detected in the paternal genome *N. tomentosiformis* and its relative *N. otophora*.

### SolRTEs mark all tobacco chromosomes, with some parent-genome specificity

To localize the physical distribution of SolRTEs, we hybridized probes of the RT and the c3’ UTR to *Nicotiana tabacum* metaphase chromosomes. Dispersed fluorescent *in situ* hybridisation (FISH) signals were detected along all chromosomes for both probes (Figure 6A), with higher abundance of the c3’ UTR-sequence, confirming computational results indicating frequent 5‘ truncation as well as the observations from Southern hybridization.

**Figure 6:**
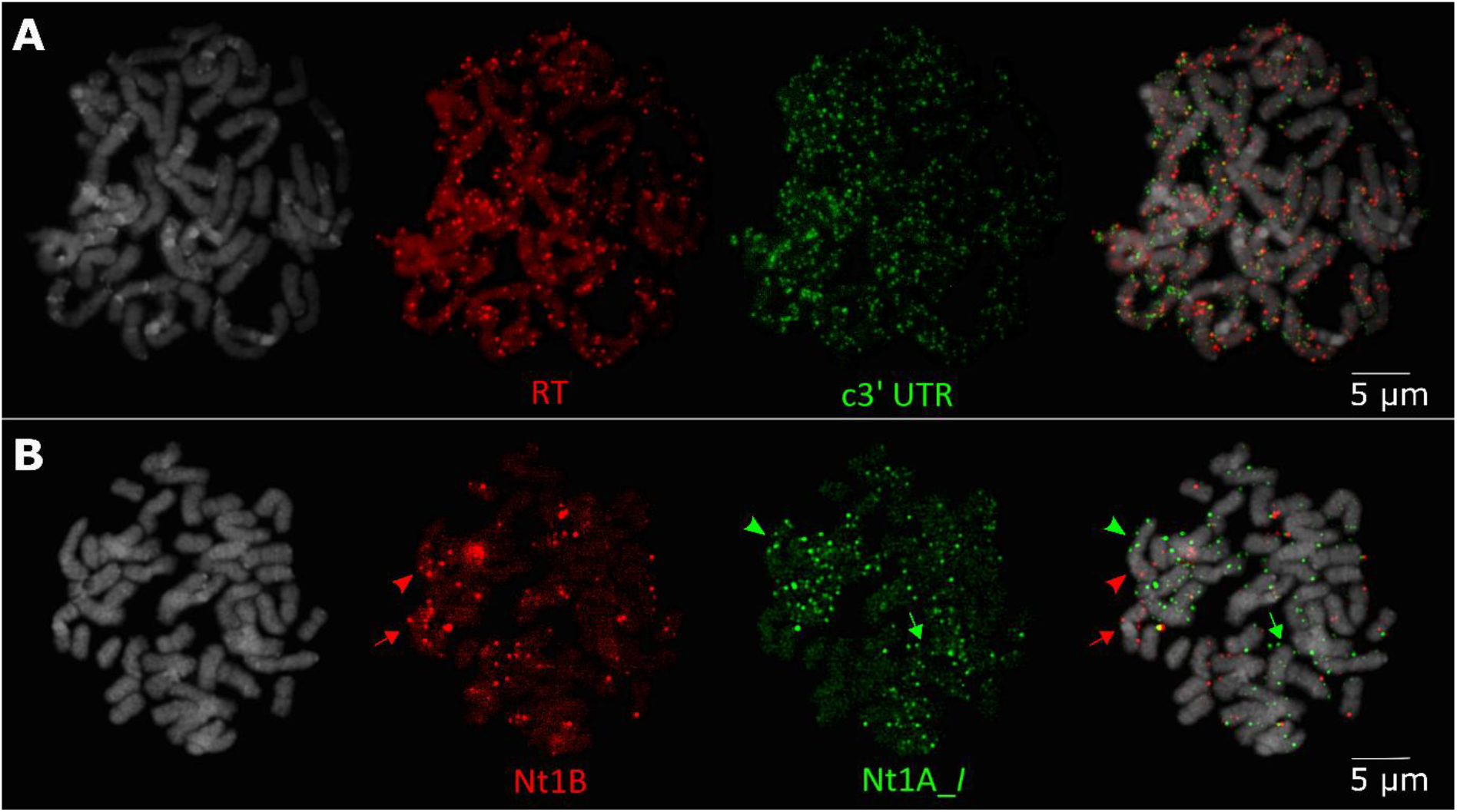
Chromosomal localisation of RTE LINEs and parental-specific SolRTE variants along *N. tabacum* metaphase chromosome spreads using multicolor fluorescent *in situ* hybridization. The DNA of N. tabacum chromosomes (grey) is stained with 4’,6-diamidino-2-phenylindole (DAPI). SolRTE LINE derived probes were either coupled to biotin (red) and digoxigenin or directly stained by DY-547 (green). (A) Diffuse signals of the reverse transcriptase domain 9 (RT, red) and conserved 3’ UTR (c3’ UTR, digoxigenin, green) showing dispersed chromosomal distribution of RTE LINEs. (B) Discrete signals of the 5’ UTRs of parental specific SolRTE lineages Nt1_B (red, *N. sylvestris* specific) and Nt1A_*I* (green, DY-547, *N. tomentosiformis* specific) show separated localisation along *N. tabacum* chromosomes. The arrows indicate examples of *N. sylvestris* (red) and *N. tomentosiformis* (green) derived chromosomes, while the arrow heads indicate an example of a recombinant chromosome.

Using 5’ UTR-derived probes of SolRTE_Nt1A_*I* and SolRTE_Nt1B, we revealed a mainly distinct localization of the parent-specific variants on tobacco metaphase chromosomes. We found chromosomes showing signals exclusively for maternal (red arrows, Figure 6B) or paternal (green arrows, Figure 6B) probes, as well as potentially recombinant chromosomes (arrowheads, Figure 6B), showing signals for both parent-specific variants, but on separated chromosome regions.

### Few SolRTE copies have potential for activation

Out of the 458 tobacco full-length SolRTEs, only four copies harboured a potentially intact, continuous ORF. All of them were assigned to the subfamily SolRTE_Nt1B, indicating this subfamily as a potentially active one within the genome of *N. tabacum* (Figure 1B, blue dots). Indeed, a copy of the SolRTE_N1B subfamily was detected in the transcriptome of *N. tabacum*. Although this subfamily was only detected within four out of the seven analyzed *Nicotiana* species, namely *N. benthamiana*, *N. attenuata*, *N. sylvestris*, and *N. tabacum*, it comprises 261 copies over all *Nicotiana* species and represents the second largest subfamily (Figure 1B & 4). Consequently, if considering the hybrid origin of *N. tabacum*, this SolRTE subfamily was most likely introduced by the maternal species *N. sylvestris*.

If the RTE subfamily SolRTE_Nt1B had been active after the hybridization event of *N. sylvestris* and *N. tomentosiformis* leading to the emergence of *N. tabacum*, empty insertion sites should exist within orthologous regions in the parental genome of *N. sylvestris*. In order to test this hypothesis, we used BLASTn similarity searches to scan the genome of *N. sylvestris* for orthologous regions matching the 53 SolRTE_Nt1B RTE insertions sites present in *N. tabacum*. Indeed, we have identified a recent SolRTE_Nt1B RTE integration, harbouring an intact and continuous ORF in the genome of *N. tabacum* contrasting the empty orthologous site in *N. sylvestris* (Figure 7). This likely indicates a recent amplification of the subfamily SolRTE_Nt1B after the formation of the hybrid species *N. tabacum*.

**Figure 7:**
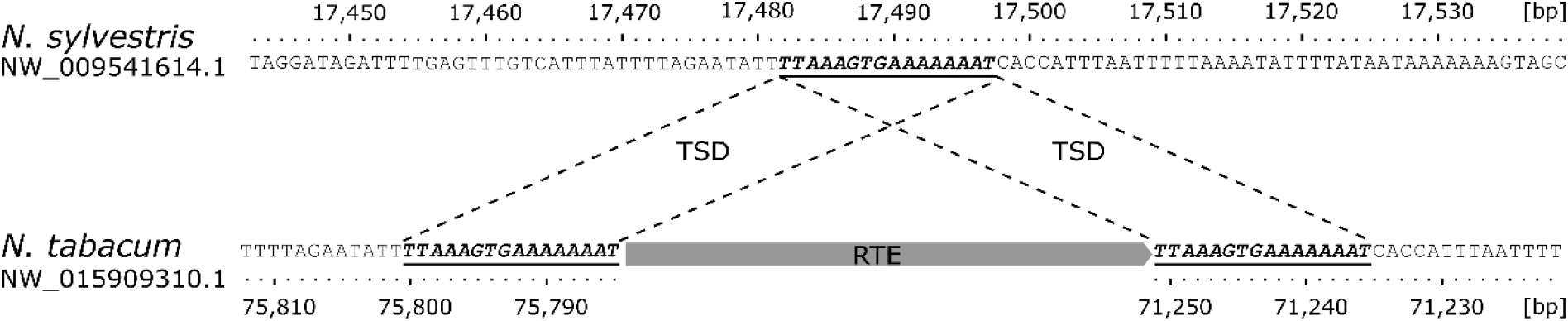
Insertion site of a continuous SolRTE_Nt1B copy in *N. tabacum*. (on contig NW_015909310.1) compared to the empty orthologous site in *N. sylvestris* (on contig NW_009541614.1). Underlined sequence was duplicated after the integration into tobacco’s genome, forming the target site duplication (TSD).

## Discussion

### UTR sequences and lengths clearly distinguish SolRTE subfamilies and variants

To provide a basis for the understanding of LINE biology and evolution in plants, we have deeply characterized the *Nicotiana* SolRTEs – a group of plant LINEs that stands out in terms of abundance, structural completeness, and regulatory potential (Renny-Byfield et al. 2011; Wenke et al. 2011; Sierro et al. 2013, 2018; Heitkam et al. 2014; Mhiri et al. 2019). Using *N. tabacum* as reference species, we provide a comprehensive overview on their sequence and structural diversity, suggesting a division into four families, eleven subfamilies, and numerous variants. Further, we traced SolRTE diversification following the speciation events within the *Nicotiana* genus, and suggest an evolutionary framework to explain the extant SolRTE diversity in nine analyzed *Nicotiana* genomes.

As most genome assemblies, especially for non-model organisms, are still incomplete and lack many repetitive regions (Peona et al. 2018) our copy number estimates are likely underrepresentations. In addition, if LINE retrotransposons are diverged or truncated, they may have escaped our computational detection or were discarded during our stringent filtering. Due to the low processivity of the RT and the potential for microhomology-mediated integration, LINEs are especially prone to 5’ truncation, as reported for mammalian and plant genomes (Noma et al. 2000; Symer et al. 2002; Komatsu et al. 2003; Zingler et al. 2005; Heitkam and Schmidt 2009). More specifically, analyzing all of the detected 4,811 SolRTE hits in *Nicotiana tabacum*, approximately 70 % were 5’ truncated. Only 10 % (458) harbored a potentially complete ORF, including the first domain of the AP-EN and the last domain of the RT. These likely include truncated copies, yet, due to the high variability of the UTRs, truncations upstream of the ORF cannot be quantified unambiguously.

In contrast to plant LINEs, their mammalian counterparts have reached much higher copy numbers, with tenthousands of similar elements within a single genome. If these mammalian L1 LINEs share an identical promoter in their 5’ UTRs, they are summarized as an L1 family (Sookdeo et al. 2013, 2018). In plants, however, the low LINE copy number is preventing a similar classification (Noma et al. 2000; Heitkam et al. 2014). Instead, we classify the SolRTE population of *Nicotiana tabacum* into the four families SolRTE_Nt1 to SolRTE_Nt4 based on the sequence similarity of their ORFs. As further subdivision into subfamilies and variants is based on the different 3’ UTRs, we classify SolRTE LINEs into at least eleven subfamilies. The 3’ UTR has a bipartite nature, being composed of a variable (v3’ UTR) and a conserved region (c3’ UTR). Each subfamily can be distinguished by their individual v3’ UTR that is guiding subfamily assignment. In contrast, for the c3’ UTR a high conservation (> 70 % nucleotide identity) across all analyzed SolRTE copies within the *Solanaceae* is evidenced consistently using bioinformatics, Southern hybridizations, and PCR experiments. The high conservation of this region is comparable to the conservation of the coding region, indicating a similar selection pressure. This high sequence conservation within the c3’ UTR may likely play a role in initiating target-primed reverse transcription as well as may serve as a recognition site for LINE-SINE partnerships (e.g. Baucom et al., 2009; Wenke et al., 2011). As a conserved 3’ terminus may be needed for mRNA recognition by the reverse transcriptase (Ohshima et al. 1996; Okada and Hamada 1997), likely by formation of a hairpin structure (Nishiyama and Ohshima 2018), we suggest that the c3’ UTR may have acquired this function.

The 5’ UTRs are even more diverse and allowed us to subclassify the subfamilies into variants. Exemplarily, we focused on the SolRTE_Nt1A subfamily that contained 111 LINE copies representing nine distinct LINE variants. Interestingly, the variability of the 5’ UTR increases towards the distal part, and hardly any copies were found to resemble each other in their full length.

### 5’ end switching as evolutionary mode for SolRTE diversification in *Nicotiana*

To trace the emergence of the SolRTE families, subfamilies, and variants, we integrate our data on SolRTE phylogenetic relationships (Figure 1) and genomic distribution (Figures 4 & 5) with the proposed *Nicotiana* phylogeny (Leitch et al. 2008; Särkinen et al. 2013). As all four SolRTE families were present across all analyzed species from the *Nicotiana* genus, but absent from any other *Solanoideae*, we propose the emergence of all four SolRTE families in the *Nicotiana* precursor (Figure 8A). This dates their emergence anywhere after the split of the so called “x=12 clade” about 24 mya (Clarkson et al. 2017; Sierro and Ivanov 2020).

**Figure 8:**
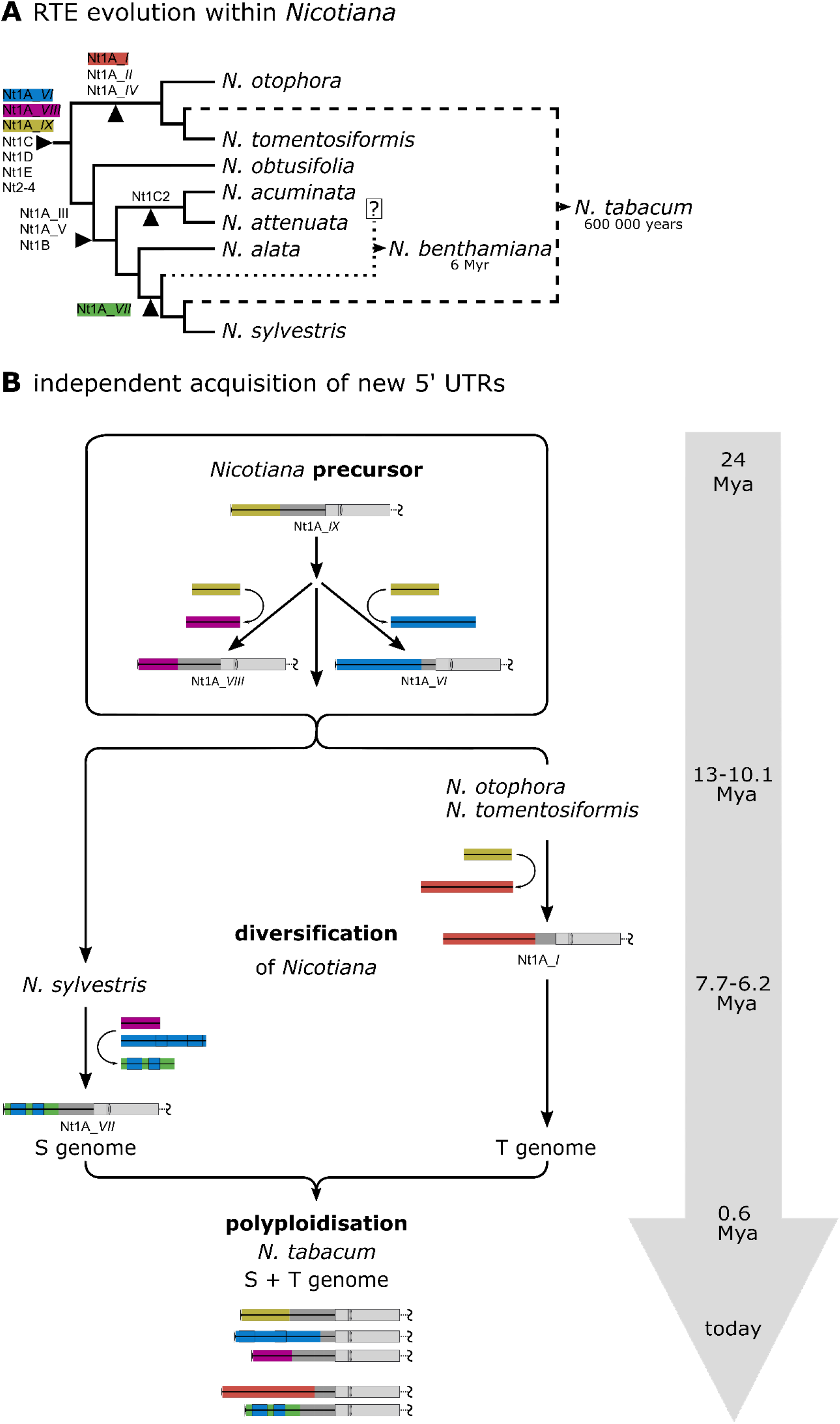
Evolutionary scenario illustrating the formation of new SolRTE variants by acquisition of non-homolgous 5’ UTRs during the diversification of *Nicotiana*. The color code corresponds to that introduced in Figure 1, highlighting SolRTE variants of interest. **(A)** Emergence of new SolRTE subfamilies and variants during the speciation of the genus *Nicotiana*. Because some lineages are distributed across all *Nicotiana* species analyzed (SolRTE_Nt2-4, SolRTENt1C-E, SolRTE_Nt1A_*VI/VII/IX*), they are likely older than the genus. Other lineages are only present in the genome of certain species, indicating the formation of new subfamilies and variants during the diversification of *Nicotiana* (e.g. SolRTENt1C2, SolRTE_Nt1B, SolRTE_Nt1A_*VII* as well as SolRTE_Nt1A). The dendrogram of the genus was adapted from Leitch *et al*., 2008) and Särkinen *et al*. (2013). **(B)** Independent acquisition of non-homologous 5’ UTR of parental specific SolRTE_Nt1A variants. Based on distribution, structural features and relationship among SolRTE_Nt1A variants we propose the following scenario: the variant *IX* is the oldest of the subfamily SolRTE_Nt1A. Within the **precursor’s** genome, the variant *IX* was amplified, but parts of the 5’ UTRs were exchanged and the variants *VI* and *VIII* emerged. During **diversification** of *Nicotiana*, the parental genomes of *N. tabacum*, e.g. ***N. sylvestris*** and ***N. tomentosiformis***, evolved independently and species-specific variants appeared. Whereas the paternal variant *I* (**T genome**) was simply formed via 5’ UTR exchange, the formation of the maternal variant *VII* (**S genome**) is more complex and included recombination among non-homolog 5’ UTRs. After the **polyploidisation** forming *N. tabacum*,parental specific variants were unity in one genome. Timeframe of evolution was adjusted according to Särkinen *et al*., 2013 and Clarkson *et al*. 2017.

To understand how plant LINE variants arise and decay, we exemplarily focus on the SolRTE_Nt1A subfamily. It contains nine variants that presently occupy the tobacco genome, all characterized by their unique 5’ UTR (Figure 2). Using the phylogenetic information from bioinformatic analyses and experiments we positioned all SolRTE_Nt1A variants across the *Nicotiana* phylogeny (Figure 8A). This enabled us to develop an evolutionary scenario explaining the observed SolRTE landscape (Figures 8B).

We trace their fates along the *N. tabacum* evolution, starting from a *Nicotiana* precursor over the diversification of the parental species into S (*N. sylvestris*) and T (*N. tomentosiformis*) genomes, to finally culminate in *N. tabacum’s* allopolyploidization. Our main conclusions are as follows:

First, we propose that variant *IX* emerged first and is the oldest of subfamily SolRTE_Nt1A (Figure 1, Figure 4, Figure 8 yellow). This is supported by our bioinformatics analyses, including ML topologies and structural comparisons.

Second, with ongoing *Nicotiana* diversification, there was continuous activity of the SolRTE_Nt1A subfamily and new variants emerged that were even section- and species-specific. This includes the variants *I*, *II*, and *IV* that arose in the T genomes of the section *Tomentosae*, as well as the variant *VII* that arose in the S genomes, exemplified by *N. sylvestris* (Figure 8B). This is supported by the SolRTE relationships (Figure 1), their presence in the respective genome assemblies (Figure 4), as well as PCR (Figure 5C,D) and FISH experiments (Figure 6).

Third, across our evolutionary scheme, we have observed at least six 5’ end switching events (Figure 8B):

i. The genome of the *Nicotiana* precursor likely already housed the three variants *VI, VIII*, and *IX* (Figure 8, blue, violet, yellow). Taking into account the ML analyses and structural element characteristics (Figures 1, 2), we hypothesize that the two variants *VI* and *VIII* emerged independently from variant *IX*, all carrying non-homologous 5’ UTRs (Figure 8B, blue, violet, yellow).
ii. A similar exchange of the 5’ UTRs likely occurred after the emergence of the T genomes, in which the variants *I, II and IV* arose (exemplarily represented by variant *I* in Figure 8B, red). Based on overall structure and phylogenetic analyses of the coding sequences, we assume that variant *IX* (yellow) has been the precursor of these variants (Figures 1, 2, 8B).
iii. The emergence of the chimeric variant *VII* in the S genome is more complex, and possibly involved several recombination events (Figure Bb, green): On one hand we observed shared motifs between the variants *VI* and *VII* (Figure 2, 8B, blue and green). These highly identical regions were embedded into otherwise dissimilar 5’ UTRs, pointing to a common origin. On the other hand we also observed a shared deletion within the v3’ UTR between the variants *VII* and *VIII* (Figure 2, 8B, green and violet). Phylogenetic analyses based on the coding sequence propose a paraphyletic relationship between the S genome specific variant *VII* and the ancient variant *VIII* (Figure 1). Therefore, we propose a chimeric structure of the variant *VII*: the main body (part of the 5’ UTR, ORF and 3’ UTR) evolved from an ancient copy of variant *VIII* (Figure 8, violet), which underwent an exchange of the 5’ UTR with a variant *VII*-specific sequence of unknown origin (Figure 8, green) and a subsequently integration of variant *VI*-derived 5’ UTR sequences (Figure 8, blue) into the otherwise non-homologous 5’ UTR of the variant *VII*.

Many different TEs contain mosaic structures as the ones observed here, allowing them to generate new and to recapture the transposable capacity of old and inactive families (e.g. (Marco and Marín 2008; Wollrab et al. 2012; Kögler et al. 2020; Maiwald et al. 2021). RNA template switching of the reverse transcriptase can be a source of new recombinant LINE copies (Bibillo and Eickbush, 2004; Gogvadze et al., 2007). If this mechanism took place between two simultaneously active LINEs, recombinated TEs such as tobacco SolRTE variant *VII* can be generated. Another source would be the 5’ fusion of a host mRNA to a LINE body. These fusion events have been already observed in fungal, mammalian and human genomes (Buzdin et al. 2002; Gogvadze and Buzdin 2005; Buzdin et al. 2007). Recruiting the promoter from a host mRNA could be advantageous, as it could provide a new transcriptionally active promoter that, if the mRNA is esssential, is unlikely to be immediately epigenetically silenced, making this a likely option for the acquisition of 5’ ends.

Alternatively, truncated LINE insertions adjacent to promoter motifs may have facilitated the acquisition of a new 5’ UTR as also has been proposed for the emergence of the composite tobacco TS-SINE (Wenke et al. 2011) and other plant LINEs (Heitkam and Schmidt 2009) and is fuelled by the high proportion of 5‘ truncated LINEs.

Anyhow, we suspect 5’ end switching as a key driver for SolRTE diversification in *Nicotiana*. As the recruitment of new 5’ ends likely goes in hand with the acquisition of new promoter regions, this strategy may be successful (i) to escape the epigenetic silencing of the host (Adey et al. 1994; Ohta et al. 2002; Khan et al. 2006; Sookdeo et al. 2013, 2018); (ii) to allow parallel activity of different LINE families within a single genome (Khan et al. 2006; Sookdeo et al. 2013, 2018); and (iii) to avoid competition for transcription factors among each other. In consequence, the high turn-over of the promoters may have rejuvenated the SolRTE landscape, secured the persistence of these LINE families in the *Nicotiana*, and likely helped them to reach comparatively high abundances.

### LINEs of the SolRTE_Nt1B subfamily are the most intact in *N. tabacum*, with limited activity after allopolyploidization

Having traced the SolRTE LINEs along the evolutionary history of *N. tabacum*, we asked, if there was any trace of LINE activity after the allotetraploidization event. This has been motivated by many reports stating that polyploidization may come along with transpositional bursts (e.g. reviewed in Vicient and Casacuberta 2017). For this, we focused on SolRTEs that are restricted to either S- or T- genomes and that are putatively active.

For both parental genomes, species-specific SolRTE variants were detected, as shown by bioinformatics and verified by PCR (Figures 4, 5). Fluorescent *in situ* hybridization of selected parent-specific SolRTE variants (SolRTE_Nt1B, SolRTE_Nt1A*_I*) to *N. tabacum* chromosomes also showed their location on S- and T-derived chromosome regions, including a few cases of co-occurrence on the same chromosome. This hybridization pattern likely reflects the contribution of each subgenome to the chromosomal landscape of *N. tabacum*.

To better understand, which SolRTEs carry the most potential for transpositional activity, we investigated their structural integrity in *N. tabacum*. We found that most ORFs were interrupted by stop codons or frameshifts, marking them as inactive and incapable for transposition. Only four copies were detected that showed continuous ORFs, all of them members of the S genome-specific subfamily SolRTE_Nt1B. This subfamily is characterized by short branch lengths in the dendrogram (Figure 1), indicating recent transposition activity. Indeed, we have identified a SolRTE_Nt1B integration site, harboring an intact and continuous ORF, in the genome of *N. tabacum* with an orthologous empty site in *N. sylvestris*. This SolRTE polymorphism identified in *N. tabacum*, but not the corresponding parental reference genome may be explained by three possibilities:

First, sequence variation among individuals of *N. sylvestris* may explain the SolRTE polymorphism. As only one genome of *N. sylvestris* is sequenced, it cannot be ruled out that another individual carries the exact TE integration. Further, incomplete lineage sorting could have resulted in removal of this integration event. Only the investigation of a large genotype panel would bring clarity.

Second, the SolRTE integration had been present in the precursor genome, but was later removed, e.g. by recombination. However, as no transposition footprints, doubled target site duplications, or similar, were detected, we consider this as unlikely.

Third, the detected SolRTE polymorphism may hint at a real, new insertion of a SolRTE that has transposed after allotetraploidization of *N. tabacum*.

Nevertheless, SolRTE_Nt1B is more abundant in *N. sylvestris* than in *N. tabacum*, and only a single polymorphism was detected between orthologous regions in *N. sylvestris* and *N. tabacum*. This suggests that SolRTE_Nt1B’s activity after allopolyploidization was limited, if present. Interestingly, SolRTE’s non-autonomous partner TS SINE (Wenke et al. 2011) showed recent activity in the polyploid tobacco and almost increased its copy number by 40 % compared to the parental genomes (Renny-Byfield et al. 2011; Mhiri et al. 2019). As the aforementioned subfamily SolRTE_Nt1B, TS SINEs are only detected within the genome of the maternal progenitor *N. sylvestris*, but not in the paternal genome *N. tomentosiformis*. As SolRTE_Nt1B is the only subfamily within the SolRTEs that may have maintained potentially active LINEs, we propose that the TS-SINE’s activity in *N. tabacum* was driven by this subgenome-specific LINE subfamily.

### Conclusion

Taken together, our in-depth SolRTE characterization across the Nightshades and especially the *Nicotiana* genus reveals how RTE LINEs evolve during the speciation and hybridization events that characterize this plant taxon. Some members, such as the SolRTE_Nt1B subfamily, were able to retain their transposition ability and showed limited activity after allopolyploidization of *N. tabacum*. Of course, our collection of full-length SolRTEs in the Nightshades is representative and can be used to annotate and mask SolRTEs in any Nightshade genome. Finally, we reveal the evolutionary strategy of RTE LINEs that allows them to keep a highly conserved LINE body, also mirrored by a second highly conserved domain in the 3’ UTR, which we name here as c3’ UTR. Indeed, this strong sequence conservation is contrasted with a highly variable 5’ UTR that is constantly renewed, and hence, rejuvenates the SolRTEs. Like this, SolRTEs likely avoid competition for transcription factors and escape epigenetic silencing. Hence, the RTE survival strategy across all Nightshades – and likely even across flowering plants – relies on pairing an evolutionary successful LINE body with a fresh 5’ UTR.

## Methods

### Identification of RTEs within genome

To detect RTE candidates, we collected the 130 reverse transcriptase domains that are representative for plant RTEs from Heitkam et al. (2014). These were complemented with 71 RTE endonucleases from a range of published elements (Heitkam et al. 2014). We aligned the respective nucleic acid sequences using MAFFT (Katoh and Standley 2013), followed by manual refinement to build nucleotide Hidden Markov Models (nHMMs). Using these nHMMs with *nhmmer* (Wheeler and Eddy 2013), we have queried the tobacco reference genome for RTE LINEs. The nHMMs and the underlying alignments are stored at https://doi.org/10.5281/zenodo.7256660.

Identified RTE candidate sequences were extracted including flanking regions of 5000 bp to each side of the RT. To filter high-confidence candidate sequences, we initially mapped all RTE-sequences against the coding region of the SolRTE-I_St1 sequence published by Wenke et al. 2011, using mapping tool in Geneious v6.1.8 (http://www.geneious.com; Kearse et al. 2012) with default settings (Table S3). From the mapped candidates, a genome-specific element was selected based on the quaility of the ORF and the presence of the first domain of the apurinic-andonuclease (Figure 1A, EN1), the ninth RT domain (Figure 1A, RT9) and the c3’ UTR (Figure 1A, c3’ UTR) used as a new reference sequence for a second mapping iteration. Both, 5’ truncated and 3’ truncated RTE copies, elements with N stretches or artificial duplicates were excluded manually from the final dataset. In particular, full-length elements were required to include the first domain of the apurinic-andonuclease (Figure 1A, EN1) and the ninth RT domain (Figure 1A, RT9). The nucleotid sequence of full-length copies were aligned using MUSCLE (Edgar 2004) followed by manual refinement in Geneious and may contain continuous or discontinuous ORFs, frameshifts or internal stop codons. Analyses of further genomes (Table S4) were carried out using the same procedure, with the pre-selected element of *N. tabacum* serving as reference for mapping.

### Classification of RTEs within *N. tabacum*

Sequence variability among putative full-length copies suggested a high diversity of *N. tabacum* RTEs. To evaluate the number of clades within the dataset, we performed a combined approach, including cluster and maximum likelihood analyses. A nucleotide sequence alignment of the ORF, including five amino acids upstream and downstream of first domain of EN and last of RT, respectively, was used. The final multiple nucleotide alignment included 458 elements and spanned 4447 bp (stored at https://doi.org/10.5281/zenodo.7256660).

For the Maximum Likelihood strategy, PartitionFinder2 (Lanfear et al. 2017) proposed a general time-reversible substitution model (GTR+G+I) as best fitting substitution model for our data. Maximum Likelihood analysis was performed using RAxML v8.2.10 (Stamatakis 2014) with 1000 bootstrap replicates. The topology was illustrated in Geneious v6.1.8 (Kearse et al. 2012).

The clustering initial into main groups the R statistical software, version 3.5.2 (R Core Team 2019) was used. Genetic distances were calculated using the ‘ape’ package (Paradis and Schliep 2019). We applied the ‘TN93’ substitution model (Tamura and Nei 1993) with pairwise deletion. Next, we estimated the best suitable number of clusters by k-means clustering using the ‘elbow method’ and transferred them to a hierarchically clustered heatmap, calculated with the ‘pheatmap’ package (Kolde 2019) and using the UPGMA distance method (Sokal and Michener 1958).

Genetic distances of each cluster were extracted and their members illustrated on the RaxML tree to verify their monophyletic origin. The complete RTE sequences underlying each cluster were compared visually in a multiple sequence alignment and checked for similarities and variation within the non-coding regions. For clusters, still showing high genetic distances and non-alignable v3’ UTRs, another cluster analysis iteration was performed. The subfamily SolRTE_Nt1A was further subdivided based on the similarity of 5’ UTRs (refer to Figure S1 – S2).

To support our subfamily classification, we used a further cluster approach using the SiLiX program package (Miele et al. 2011). Clustering series were performed in which the identity threshold was gradually increased from 70 % to 100 % (5 % steps) over two sequence length thresholds (80 % and 100 %) of the ORF. A minimum cluster size of five was required. The appropriate number of clusters was detected by a sudden decrease in the total number of sequences assigned to a cluster and an increase in the number of proposed clusters (refer to Figure S3).

### Characterisation of RTEs within *Nicotiana tabacum*

The selection of reference elements for each subfamily and variant was based on three main criteria: (1) the presence of a TSD, (2) the intactness of the ORF, and (3) the full element length. TSDs were detected manually by self-dotplot comparison using the implemented tool in Geneious. As the TSD belongs to the host insertion site, it was not treated as part of the LINE copy. Start and end points of the ORFs were transferred from the SolRTE_Nt1B reference, as this is the only clade with a continuous ORF. To define the c3’ UTR, we relied on the sequence identities across an alignment of all reference elements. We defined the first base of the c3’ UTR as the position in the alignment that showed 90% identity within the 3’ UTR and was followed by a conserved sequence. The RTE LINE ends before the first TTG-triplett. Only complete tripletts were counted for the TTG-tail and up to one point mutations were tolerated. Variable 5’ UTR and 3’ UTR were defined as regions between the upstream TSD and the ORF and between the ORF and the c3’ UTR, respectively. The 5’ UTR sequence of each reference element was checked for promoter motifs using the PlantCare database (Lescot et al. 2002). The Alignment of the 19 representative SolRTE LINEs of tobacco were stored at https://doi.org/10.5281/zenodo. 7256660.

### Relationships of Nightshade RTEs

To examine the relationships of SolRTEs in Nightshade species, k-means and hierarchical clustering were performed as described above. 10 % or at least 20 elements were randomly selected as representative RTE full-length copies of each genome. For *S. tuberosum* only eight full-length elements were detetcted and all were included. For *S. lycopersicum* no full-length element was detected, therefore this species was completely excluded.

### Distribution of RTEs within the genus *Nicotiana*

To investigate the distribution of the identified RTE lineages within the genus *Nicotiana*, we mapped each species-specific SolRTE population independently against each reference element of *N. tabacum* using the implemented tool in Geneious (Table S3). Species-specific Neighbour Joining trees were calculated based on the RTE nucleotide alignment mentioned above and under the assumption of the ‘Tamura-Nei’ substitution model (Tamura and Nei 1993). All members of a monophyletic clade were counted as member of the according lineage. The distribution was illustrated using the R package ‘ggplot2’ (Wickham 2016).

### Extracting empty sites of RTE insertion in parental species and potentially active RTE elements

All SolRTE_Nt1B members from *N. tabacum* and *N. sylvestris* were mapped against each other including their flanking regions using the Geneious mapper tool (Table S3). Out of 53 SolRTE_Nt1B copies of *N. tabacum* 28 were mapped against *N. sylvestris*, leaving 25 potential new insertions sites. The RTE copies themselves were cut out to create artifical empty insertion sites used for the identification of an orthologous site within the maternal genome *N. sylvestris* using BLASTn similarity searches in Geneious. Members of the SolRTE_Nt1B subfamily containing continuous and complete ORFs were used to detect transcriptionally active copies using online BLASTn similarity searches on NCBI within the ‘transcriptome shotgun assembly’ of *N. tabacum*.

### Plants and DNA isolation

Seeds of the following plants were obtained by the IPK Gatersleben and grown under long-day conditions (16h, 25°C) in a greenhouse: *Nicotiana tabacum* ‘Rote Front’, *N. tomentosiformis* ‘Goodspeed’ (NIC 479), *N. sylvestris* (NIC 6), *N. attenuata* (PI 555476), *N. benthamiana ‘*Domin*’* (NIC 660), *N. otophora* (PI 302477), *N. acuminata* (NIC 471), *N. alata* (NIC 431), *Solanum tuberosum* ‘Gala’, *S. lycopersicum* ‘Sparta’*, S. melongena (*SOL 1041), *Capsicum annuum* ‘Paprika de Cayenne’*, Petunia axillaris* (PETU 2), *Physalis peruviana* (PHY 36), *Ipomoeae purpurea*(IPO 5), *Ocimum basilicum var. basilicum* (OCI 370). Young leaves were collected for genomic DNA isolation using the cetyl-trimethyl/ammonium bromide (CTAB) protocol (Saghai-Maroof et al. 1984).

### Polymerase chain reaction (PCR)

To verify the presence and distribution SolRTEs among *Solanaceae*, we designed primers to amplify specific regions of the SolRTEs (Table S5). PCR with 100 ng template DNA was performed in 50 μl volume containing 0.2 mM dNTPs, 10 pg of the primer, 1 × DreamTaq buffer, and 1 U DreamTaq (Thermo Fisher Scientific). PCR conditions were as usual: 93°C for 3 min, followed by 30 cycles of 93°C for 20 sec for denaturation, the primer-specific annealing temperature (Table S5) for 30 sec, 72°C for 30 sec, and a final incubation at 72°C for 5 min.

For hybridization experiments, PCR products were purified (As One International,Invisorb Spin DNA Extraction Kit) and ligation into pGEM-T plasmids (ThermoFisher Scientific, manufacturer’s conditions) of the PCR fragments were conducted for sequencing and probe preparation. Sequences of the probes are stored at https://doi.org/10.5281/zenodo.7256660.

### Southern hybridization

For gel blots, *Hinf*I-restricted genomic DNA was separated on a 1.2 % agarose gel and transferred onto Hybond-N+ nylon membranes (GE Healthcare) by alkaline transfer. Hybridization was performed under standard conditions and incubated at 60°C using radioactive probes labelled by random priming (Sambrook et al. 1989). Membranes were washed at 60°C for 10 min in 2 × standard saline citrate (SSC)/ 0.1 % sodium dodecyl sulfate (SDS), followed by 10 min in 1 × SSC/ 0.1 % SDS.

### Fluorescent *in situ* hybridisation

For preparation of mitotic chromosomes, young leaves were synchronized for 3 to 5 h in 2 mM 8-hydroxyquinoline, fixated in methanol:acetic acid (3:1), and macerated in an enzyme mixture containing 2.0 % (w/v) cellulase from *Aspergillus niger* (Sigma-Aldrich), 4.0 % (w/v) cellulase Onozuka-R10 (Serva), 5 % (v/v) pectinase from *A. niger* (Sigma-Aldrich), 0.5 % (w/v) pectolyase *A. niger* (Sigma-Aldrich), and 2.0 % (w/v) cytohelicase from *Helix pomatia* (Sigma-Aldrich). Subsequently, the nuclei suspension was dropped onto slides as described by Desel et al. (2001). Probes of specific SolRTE-regions were labelled indirectly by PCR in the presence of biotin-11-dUTP or digoxigenin-11-dUTP and directly in the presence of DY647-dUTPs.

Fluorescent *in situ* hybridisation (FISH) of *N. tabacum* chromosomes was performed according to Heslop-Harrison et al. (1991). Stringencies of 76 % for hybridisation and 79 % during washing steps were used. Using 4’,6-diamid-2-phenylindol (DAPI), chromosomes were counterstained and mounted in antifade solution. Slides were examined with an “Axioplan 2 Imaging” microscope (Carl Zeiss) and images were acquired using the digital FISH analysis system “GenASIs” (Applied Spectral Imaging) coupled with high-resolution CCD camera SDI BV300-20A. Images were optimized using Adobe Photoshop 5 software.

## Data access

All data to reproduce this study have been made accessible under https://doi.org/10.5281/zenodo. 7256660. This includes the nHMMs and underlying nucleotide sequence alignments, the multiple nucleotide alignment of 458 full length SolRTE LINEs, the 19 representative SolRTE LINEs of tobacco and the sequences of SolRTE LINE hybridization probes.

## Acknowledgments

We thank Ines Walter and Susan Liedtke for their excellent technical help and Dr. Beatrice Weber and Sophie Maiwald for inspiring discussions. We remember the late Prof. Dr. Thomas Schmidt, who has initialized and supported this study. The gene bank of the IPK Gatersleben is gratefully acknowledged for providing plant seeds.

