## Supplementary Material for "How to start a LINE: 5’ switching rejuvenates LINE retrotransposons in tobacco and related *Nicotiana* species"

Article acceptance date:

This PDF file includes:

Supplemental Notes S1

Supplemental Figure S1 – S4

Supplemental Table S1 – S5

Supplemental References

Notes S1: **Identification of full-length SolRTEs (Details).**

In order to identify partial and full-length RTE LINEs in *N. tabacum*, we used an nHMM algorithm for the detection of RTs of the RTE-type. We identified 7159 potential RTE hits in the genome of *N. tabacum* TN90. All in all 4811 hits have been mapped against SolRTE-I_St1 sequence published by Wenke et al. (2011), indicating them as related to this reference sequence and sharing a minimum of 250 bp. Out of these mapped candidates 3247 and 307 sequences showed 5’ truncated or 3’ truncated RTE copies, respectively and 799 sequences were excluded as they harbor long N stretches, the extracted contigs were truncated or artificial duplicates of each other. Unmapped sequences were probably too old copies and therefore too diverged or share too short regions to be compared to the reference sequence, those can be excluded of the analyses as an alignment would be impossible. In total 458 RTE elements within the genome of *N. tabacum* were defined as full length elements, with an open reading frame (ORF) spanning the first domain of the apurinic-endonuclease (Figure 1A, EN1) and the last domain of the Reverse Transcriptase (Figure 1A RT9), no matter if the ORF was continuous or discontinuous, harbors a frameshift or any internal stop codons.

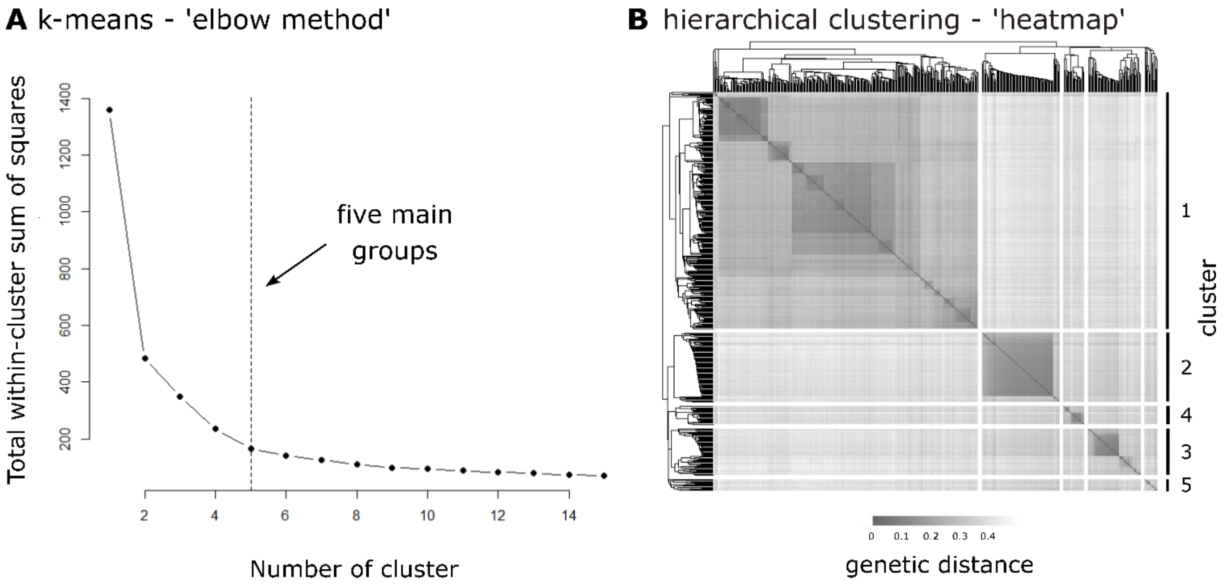


**C** inter - & intra cluster genetic distances

| cluster | 1 | | 2 | | 3 | | 4 | | 5 | |
|---|---|---|---|---|---|---|---|---|---|---|
| cluster1 | 0.27 | | | | | | | | | |
| (283) | 0.01 | 0.43 | | | | | | | | |
| cluster2 | 0.4 | | 0.16 | | | | | | | |
| (83) | 0.29 | 0.49 | 0.08 | 0.38 | | | | | | |
| cluster3 | 0.4 | | 0.35 | | 0.3 | | | | | |
| (56) | 0.29 | 0.48 | 0.27 | 0.45 | 0.07 | 0.41 | | | | |
| cluster4 | 0.38 | | 0.35 | | 0.35 | | 0.25 | | | |
| (22) | 0.27 | 0.47 | 0.27 | 0.45 | 0.27 | 0.44 | 0.09 | 0.36 | | |
| cluster5 | 0.4 | | 0.35 | | 0.35 | | 0.35 | | 0.32 | |
| (14) | 0.31 | 0.49 | 0.28 | 0.42 | 0.29 | 0.43 | 0.27 | 0.41 | 0.09 | 0.37 |

Figure S1: **Initial clustering of the 458 SolRTE identified within *N. tabacum* 'TN90' indicates five main SolRTE groups.** Using the TN93 substitution model (Tamura and Nei 1993), a genetic distance matrix was calculated for the conserved nucleotide sequences that span the open reading frame from the first apurinic endonuclease to the last reverse transcriptase protein domains and used for cluster methods. **(A)** To estimate an appropriate number of clusters within the dataset, we used the 'elbow method'. This calculates the number of clusters, which explains the majority of variance within a given dataset. The dashed line indicates the determined number of clusters. **(B)** The determined number of clusters (5) was transformed to a hierarchical clustered heatmap. Intensity of the grey shading corresponds to the genetic distance. Dendrograms and clustering were based on UPGMA distance method. **(C)** Genetic distances within and among clusters of SolRTEs. Mean genetic distances and the span of the minimum and maximum genetic distance were shown. Number of RTE LINEs of each cluster are mentioned in brackets.

### Genetic distances within and between SolRTE main groups

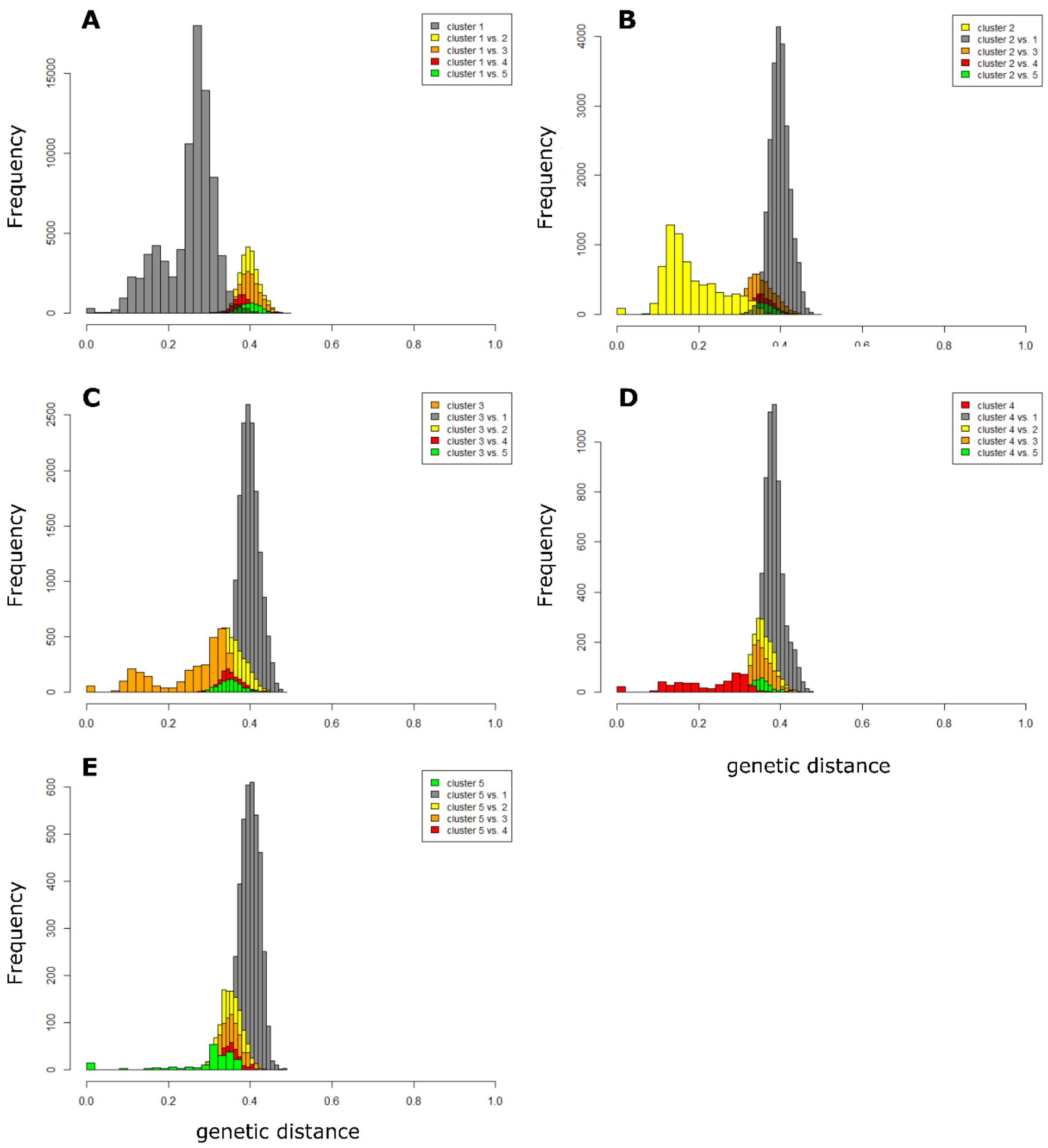


Figure S2: **Bar plots of the genetic distance within and among each determined cluster of the 458 SolRTE identified within *N. tabacum* 'TN90'.** Intra genetic distances are indicated for **(A)** cluster 1 in grey, **(B)** cluster 2 in yellow, **(C)** cluster 3 in orange, **(D)** cluster 4 in red and **(E)** cluster 5 in green. Inter genetic distances to each cluster are indicated in the affiliated color.

**Fine clustering supports the SolRTE classification into subfamilies**

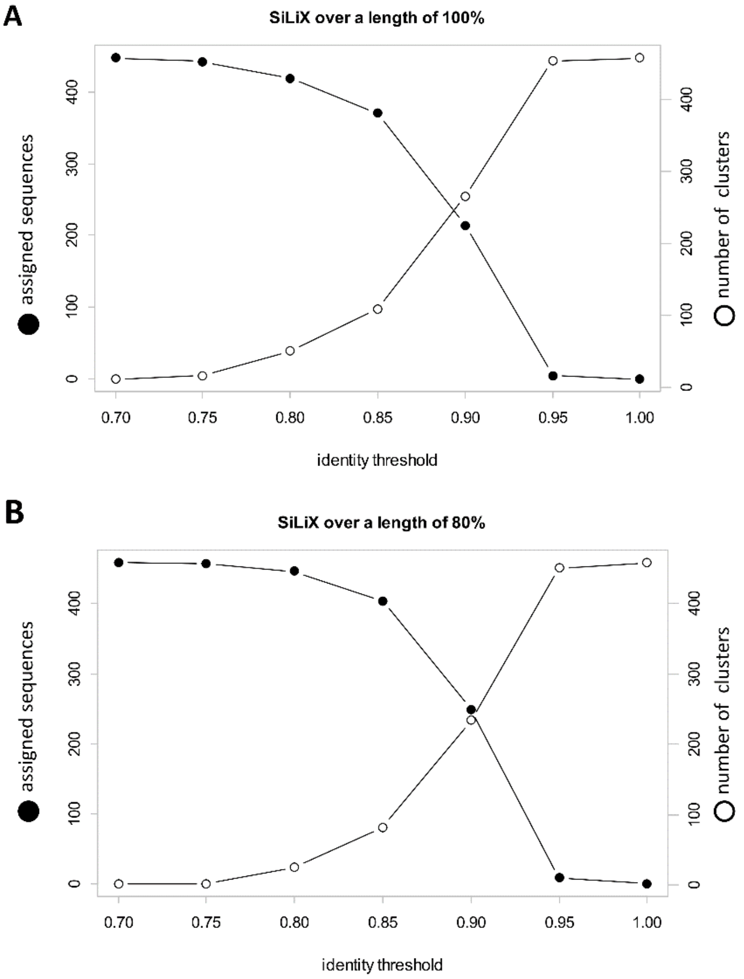


Figure S3: **SiLiX cluster analysis of 458 SolRTE members identified in the genome of *N. tabacum* 'TN90'**. SiLiX cluster analysis assigns sequences into groups based on predetermined identity thresholds within one cluster over a given length. To determine an appropriate number of cluster within the SolRTE dataset of *N. tabacum*, two SiLiX clustering series were performed. For this purpose, the identity threshold was increased stepwise (5%) from 70 % to 100 % over a sequence length of 80 % and 100 %, respectively. The appropriate number of clusters were determined by a strong decrease in the number assigned to a cluster and a strong increase in the number of proposed clusters. For both length parameters, the number of sequences assigned to a cluster decreased sharply and the number of proposed clusters increased sharply by raising the identity threshold from 85 % to 90 %. With an identity threshold of 85 %, SiLiX cluster proposed 9 or 8 cluster with more than 5 members at lengths of 80 % and 100 %, respectively. These clusters support the subfamily structure determined by k-means and hierarchical clustering. Compared to the first two methods, SiLiX estimated smaller cluster sizes and excluded many sequences based on the applied length and identity thresholds. The SolRTE_Nt2A, SolRTE_Nt3A and SolRTE_Nt4A subfamilies were excluded completely. On the other hand, SiLix cluster analysis divided subfamily SolRTE_Nt1B into two clusters.

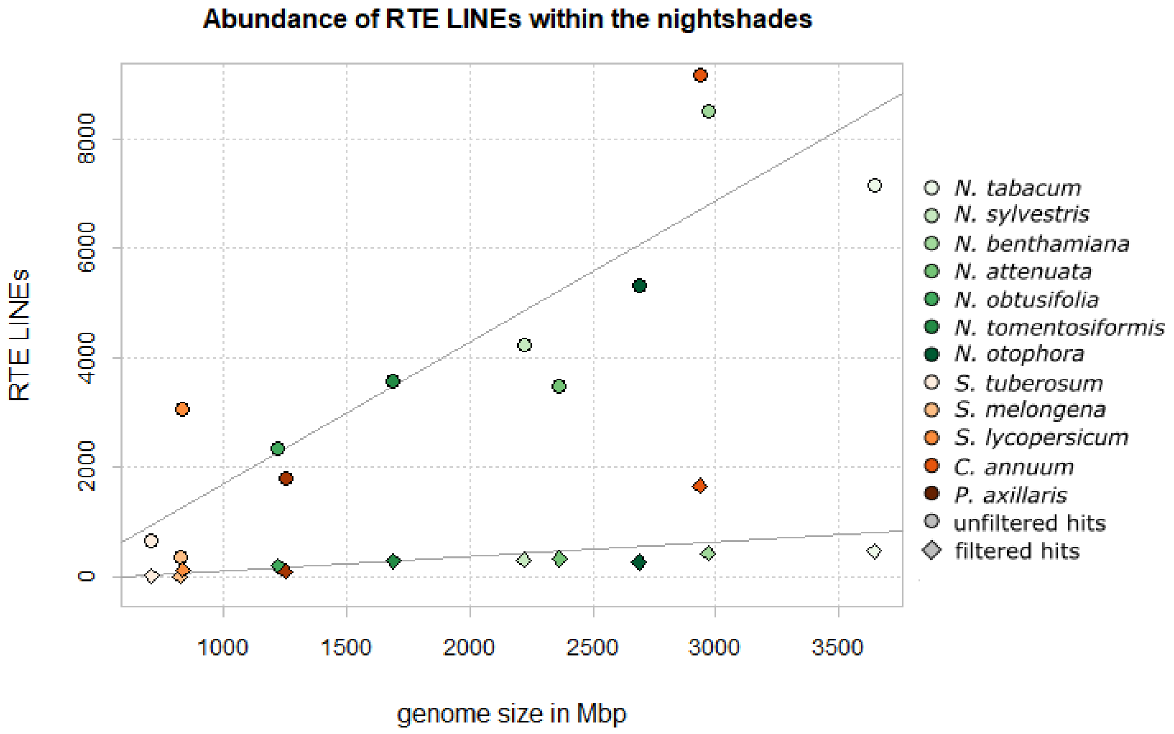


Figure S4: **Abundance of RTE LINEs within the nightshades (including the genera *Nicotiana*, *Solanum*, *Capsicum* and *Petunia*) in correlation to their genome size in Mbp**. Number of unfiltered hits were taken directly from the nHMM output data and are depicted as rectangles. After manual inspection, the number of filtered hits included RTE LINE copies with open reading frames spanning from the first apurinic endonuclease to the last reverse transcriptase protein domains. Filtered hits were depicted as circles.

Table S1: **Lengths of representative RTE LINEs and their respective structural features of each identified SolRTE family (Nt1-4), subfamily (marked by the appendices A-E) and the nine variants of the subfamily SolRTE_Nt1A (SolRTE_Nt1A_*I* – *IX*) in *N. tabacum* 'TN90'**. Length of full LINEs, their open reading frames (ORF), untranslated regions at the 5' and 3' terminus (5' UTR and 3' UTR, respectively), the conserved 3' UTRs (c3' UTR) and target site duplications (TSD) are denoted in base pairs (bp), whereas the repetitive motif at the 3' terminus ([TTG]n) is counted as numbers of repetitions. We observed variable element lengths, ranging between 3865 bp and 4718 bp. As the length of the ORF (2869 – 3021 bp, median 2964 bp) and the c3' UTR (95 – 106 bp, median 104 bp) is conserved over all reference elements, the variable full length of an element reflects the sizes of its highly variable and non-homologous 5' UTR (534 – 1384 bp, median 872 bp) and its v3' UTR (144 – 314 bp, median 232 bp). *N. tabacum* RTE LINEs are 3' terminated by a sequence of [TTG]-tripletts with 4 up to 23 repetitions. All reference elements are flanked by target site duplications (TSDs) between 10 and 21 bp in length, with a median length of 14 bp.

| RTE LINE | length (bp) | | | | | | number of repetition |
|---|---|---|---|---|---|---|---|
| | full LINE | 5' UTR | ORF | v3' UTR | c3' UTR | TSD | $[TTG]_n$ |
| SolRTE_Nt4B | 3897 | 586 | 2869 | 314 | 100 | 12 | 9 |
| SolRTE_Nt4A | 4185 | 963 | 2922 | 184 | 103 | 18 | 4 |
| SolRTE_Nt3B | 4484 | 1154 | 2938 | 213 | 105 | 11 | 23 |
| SolRTE_Nt3A | 4016 | 771 | 2953 | 166 | 104 | 15 | 7 |
| SolRTE_Nt2B | 4408 | 1164 | 2918 | 201 | 105 | 17 | 6 |
| SolRTE_Nt2A | 4520 | 1234 | 2947 | 219 | 105 | 17 | 5 |
| SolRTE_Nt1E | 4587 | 1206 | 2966 | 296 | 104 | 11 | 6 |
| SolRTE_Nt1D | 3918 | 596 | 2972 | 232 | 104 | 14 | 5 |
| SolRTE_Nt1C | 4212 | 843 | 2986 | 252 | 106 | 14 | 8 |
| SolRTE_Nt1B | 3865 | 534 | 2977 | 232 | 103 | 11 | 6 |
| SolRTE_Nt1A_*IX* | 4255 | 872 | 3021 | 228 | 95 | 10 | 14 |
| SolRTE_Nt1A_*IIX* | 4011 | 765 | 2950 | 178 | 104 | 13 | 5 |
| SolRTE_Nt1A_*VII* | 4127 | 827 | 2987 | 197 | 104 | 16 | 4 |
| SolRTE_Nt1A_*VI* | 4270 | 925 | 2964 | 257 | 103 | 15 | 7 |
| SolRTE_Nt1A_*V* | 4016 | 685 | 2974 | 235 | 104 | 10 | 7 |
| SolRTE_Nt1A_*IV* | 4718 | 1384 | 2981 | 235 | 104 | 21 | 6 |
| SolRTE_Nt1A_*III* | 3889 | 595 | 2941 | 233 | 104 | 17 | 8 |
| SolRTE_Nt1A_*II* | 4421 | 1177 | 2870 | 243 | 104 | 17 | 9 |
| SolRTE_Nt1A_*I* | 4385 | 1050 | 2979 | 243 | 104 | 12 | 7 |

Table S2: **Abundance of RTE LINEs in each analyzed nightshade genome** (including the genera *Nicotiana*, *Solanum*, *Capsicum* and *Petunia*). Number of unfiltered hits were taken directly from the nHMM output data and are depicted as rectangles. After manual inspection, the number of filtered hits included RTE LINE copies with ORF spanning from the first apurinic endonuclease (EN1) to the last reverse transcriptase protein domains (RT9). The identity and mean length of their open reading frames (ORF) were calculated from the domain EN1 up to RT9.

| Species | Unfiltered hits | Filtered hits | Mean length ORF [bp] | Identity % |
|---|---|---|---|---|
| *Nicotiana tabacum* | 7159 | 458 | 2669.1 | 71.9 |
| *Nicotiana sylvestris* | 4229 | 302 | 2666.9 | 74.6 |
| *Nicotiana tomentosiformis* | 3565 | 282 | 2671.0 | 74.1 |
| *Nicotiana attenuata* | 3469 | 319 | 2670.0 | 75.0 |
| *Nicotiana benthamiana* | 8500 | 392 | 2658.9 | 69.6 |
| *Nicotiana obtusifolia* | 2329 | 184 | 2654.9 | 72.2 |
| *Nicotiana otophora* | 5317 | 247 | 2668.8 | 70.7 |
| *Solanum tuberosum* | 663 | 8 | 2601.0 | 76.3 |
| *Solanum melongena* | 3043 | 109 | 2688.6 | 73.4 |
| *Solanum lycopersicum* | 352 | -- | -- | -- |
| *Capsicum anuum* | 9160 | 1646 | 2670.2 | 72.5 |
| *Petunia axillaris* | 1787 | 83 | 2660.5 | 73.3 |

Table S3: **Mapping parameters** used with the Geneious mapper.

| | Maximum gaps per read % | Maximum Gap Size bp | Minimum Overlap bp | Minimum Overlap Identity % | Word length | Index Word length | Maximum mismatches per read % | Maximum ambiguity |
|---|---|---|---|---|---|---|---|---|
| Identification of RTE full-length elements within genomes | 50 | 80 | 250 bp | - | 4 | 4 | 80 | 64 |
| Distribution of classified SolRTE families / subfamilies / variants within Genomes of analyzed *Nicotiana* species | 25 | 20 | length of reference element | 80 | 4 | 4 | 75 | 64 |
| Extracting empty sites of the SolRTE_Nt1B subfamily insertion in parental species | 5000 | 300 | 5000 | 80 | 4 | 4 | 90 | 64 |

Table S4: Genome assemblies and accession numbers of the analyzed nightshade species

| Species | | Source | Accession | References |
|---|---|---|---|---|
| *Capsicum* | *Anuum* | GenBank | GCF_000710875 | Qin et al. 2014 |
| *Nicotiana* | *attenuata* | GenBank | GCF_001879085.1 | Xu et al. 2017 |
| *Nicotiana* | *benthamiana* | SolGenomics | Niben.genome.v1.0.1.scaffolds.nrcontigs | Bombarely et al. 2012 |
| *Nicotiana* | *obtusifolia* | GenBank | GCA_002018475.1 | Xu et al. 2017 |
| *Nicotiana* | *Otophora* | GenbAnk | GCA_000715115 | Sierro et al. 2014 |
| *Nicotiana* | *Sylvestris* | GenBank | GCA_000393655.1 | Sierro et al. 2013 |
| *Nicotiana* | *Tabacum* | GenBank | GCF_000715135.1 | Sierro et al. 2014 |
| *Nicotiana* | *tomentosiformis* | GenBank | GCA_000390325.2 | Sierro et al. 2013 |
| *Petunia* | *Axillaris* | SolGenomics | Petunia_axillaris_v1.6.2_genome | Bombarely et al. 2016 |
| *Solanum* | *lycopersicum* | GenBank | GCA_000188115-2 | Sato et al. 2012 |
| *Solanum* | *melongena* | GenBank | GCA_000787875.1 | Hirakawa et al. 2014 |
| *Solanum* | *tuberosum* | GenBank | GCF_000226075 | Xu et al. 2011 |

Table S5: **Primer pairs** used for amplification within *N. tabacum* and other nightshades

| SolRTE family / structure | Primer | | Amplicon | Annealing temperature | study |
|---|---|---|---|---|---|
| SolRTE_c3' UTR | for | TGCTTTCTTGAGCCGAGG | 125 bp | 58.8 °C | this study |
| | rev | AACAACATACCCAGTATAATCCC | | | |
| SolRTE_v3' UTR | for | AGATGAGGATGTTGAGGTGG | 665 bp | 56°C | this study |
| | rev | AACAACATACCCAGTATAATCCC | | | |
| SolRTE-I_RT | for | AGATGAGGATGTTGAGGTGG | 235 bp | 53.8 °C | Wenke et al. 2011 |
| | rev | TCTCCCCAATACTTCTTAGG | | | |
| SolRTE_Nt2B | for | CGTTTGTGACCTTTAGTTGTAG | 300 bp | 54.8 °C | this study |
| | rev | ATTACCGAAATATAATCTACCAC | | | |
| SolRTE_Nt1B | for | TCAGGGAAATTCTCGAAGTTG | 145 bp | 54.8°C | this study |
| | rev | TGTGTTGATTTGTTGTACTCG | | | |
| SolRTE_Nt1A_*I* | for | TATTTTCTGGCGCACTGACC | 275 bp | 56.8 °C | this study |
| | rev | GAAAACTGGAAAACGGGAGG | | | |
| SolRTE_Nt1A_*VII* | for | TCCCCATCATCCAACACC | 345 bp | 58.8 °C | this study |
| | rev | CGGTCCAAACTGATACTGTC | | | |
